# Computational identification of responding T cell clones from longitudinal repertoire sequencing

**DOI:** 10.64898/2026.05.29.728700

**Authors:** Martina Milighetti, Zachary Sethna, Stephen Martis, Charlotte Reiche, Yuval Elhanati, Vinod P. Balachandran, Benjamin D. Greenbaum, Aleksandra M. Walczak, Thierry Mora

## Abstract

T cells activate and expand upon interaction with cognate antigen, derived from pathogens or mutated proteins. T cell clones can be identified by their T cell receptor (TCR) which acts as a unique barcode to track their expansion. Longitudinal TCR sequencing can be used to track T cell responses to a large array of stimuli. However, experimental identification of T cell clones of interest is challenging, especially when information about the driving antigen is lacking. Computational identification based on clonal dynamics is an antigen-agnostic alternative. However, it is subject to sequencing noise and biological variability, and relies on the choice of particular timepoints that are compared to find expanding and contracting clones. We review existing methods for identification of responding T cell clones from TCR sequencing, and the strategies that have been used to account for noise. We highlight that most methods look for clones at specific timepoints, and discuss the need for unbiased methods to shed light on new dynamics. We showcase how to address these needs by proposing CloneSearch, a method to identify expanding and contracting T cell clones from longitudinal TCR sequencing which is agnostic to the time of stimulus and can account for the noise these clones are subject to. CloneSearch can recapitulate previously identified responses from published data, and expand the analysis to show identification of previously undetected responses from these same datasets.

## I. INTRODUCTION

T cells are key effector cells of the adaptive immune system that activate and proliferate upon immune stimulation. T cells express a T cell receptor (TCR), most commonly composed of a single *α* and single *β* chain (*αβ*TCR), on their surface. The TCR mediates antigen recognition by binding to an antigen (or peptide, p) presented on a major histocompatibility complex (MHC) encoded by the human leukocyte antigen (HLA) gene in humans. TCR-pMHC binding leads to surface expression of activation markers and production of cytokines by T cells, followed by proliferation and differentiation of the cell into different effector and memory functions.

T cells need to recognise a wide variety of antigens to ensure protection against all possible pathogens. To achieve such wide recognition, the TCR needs to be very diverse [1]. Diverse TCRs are generated through somatic V(D)J recombination, a random process of DNA rearrangement that takes place in each T cell and generates a diversity of upwards of 10^23^ potential nucleotide sequences [2]. Because of this process and the diversity it can generate, the TCR repertoire, defined as the set of all TCRs expressed in an individual, can react to a large number of immune stimuli. Moreover, the generated diversity causes each T cell clone in an individual to have an essentially unique TCR nucleotide sequence. Thus, the TCR provides a natural barcode to track the expansion and contraction of single T cell clones in the repertoire, and the TCR repertoire contains information on the history of immune stimuli encountered by an individual [3].

Not all stimuli are equally immunogenic [4–7]. Some antigens can induce abundant and long-lasting T cell responses, while others activate T cells more weakly or transiently. By measuring the magnitude and duration of T cell responses over time, we can estimate immune protection against a pathogen [8–10], expected protection from vaccination [11–14], or pre-existing protection against an infection [15, 16]. This is of particular interest in the field of cancer immunotherapy, where predicted immunogenic antigens are a good predictor of survival or response to immune checkpoint blockade [5, 6, 17, 18], and are key information in the generation of cancer vaccines [12, 19–25].

Here, we provide an overview of existing methods for the estimation of T cell responses. We particularly focus on computational methods for the analysis of longitudinal TCR*β* chain sequencing datasets. We discuss the noise properties of four datasets with previously identified responses [8, 10, 13, 14] and review existing methods for the identification of responding TCRs from longitudinal datasets. We outline two issues that are not addressed in current methods: (1) the need to account for clone size-dependent noise in fold-change calculations without requiring sequencing replicates and (2) the identification of background and responding sequences in a way that is agnostic to the timing of immune stimulus. We discuss a mathematical transformation that can be used as an alternative to naïve fold changes to account for the different sources of noise dominating for small and large clones and to better estimate which clones expand (or contract) beyond the expected noise. We further highlight that existing methods rely on knowing the time of the immunological challenge and expected dynamics of T cell expansion for the identification of responding clones. We argue that unbiased identification of responding clones across the time series can further expand our understanding of T cell responses by uncovering previously unobserved dynamics in the TCR repertoire. We summarize these observations in the form of a PCA-based method which identifies a bulk of sequences that do not change over time (the “background”), as well as a handful of sequences that have unexpected temporal trajectories (the “outliers”). We make the proposed methodology available to the community through a Python package which we call CloneSearch. Using four datasets with previously identified responses, we use CloneSearch to show how unbiased approaches that pay attention to noise can successfully identify previously annotated clones without any information on when the stimulus occurred as well as pick up responses that had not been previously identified.

## II. MEASURING THE T CELL RESPONSE

When the antigen or antigens of interest are known, the magnitude and duration of the T cell response can be measured experimentally by peptide stimulation or tetramer staining experiments (fully reviewed in Refs. [26, 27]). A peptide stimulation experiment can either consist of a co-culture of T cells with antigen-pulsed antigen presenting cells or in vitro peptide stimulation of unsorted peripheral blood mononuclear cells (PBMCs), where the antigen is either a single peptide or a peptide pool. In these conditions, T cells that are specific for the pulsed antigen get activated, and will start over-expressing characteristic activation markers. Activation can be measured for the whole T cell population by ELISpot [28], or the expression levels of activation-induced markers (AIM) on single cells can be measured by flow cytometry [29–33]. Alternatively, multimers can be used to identify T cells with a TCR specific for the antigen of interest. Multimers are peptide-loaded MHC molecules which are conjugated with a fluorophore. By staining a T cell population with an MHC multimer loaded with the antigen of interest, T cells specific for the antigen can be identified [34–44]. While both AIM and pMHC multimer staining allow for sorting of single antigen-specific cells and can reliably identify specific T cells and their receptor through further sequencing, they are time-consuming and require the previous identification of immunogenic antigens to screen, which depend on each patient’s ability to present relevant peptides on their HLA.

Computational analysis of repertoire sequencing experiments can address some of these limitations. Because a TCR nucleotide sequence is unique to a T cell clone, it can be used as a barcode to identify single T cell clones by TCR sequencing [45, 46]. T cell clones of interest can then be identified through either their sequence characteristics or their dynamics over time. These approaches do not rely on knowing the antigen driving the response *a priori* and can therefore be used as a screening method to identify a T cell response against any antigen.

T cells that recognise the same antigen often show high sequence similarity [47–49]. This observation can be leveraged to identify sequences that respond to the same stimulus by looking for TCR sequences that are more similar to each other than expected by chance [50]. The sequence-based approach has proven to be a successful screening method in the identification of, among others, COVID-specific TCRs [51] or *Mycobacterium tuberculosis*-specific (Mtb-specific) TCRs [52]. However, TCR sequence similarity is neither necessary nor sufficient for shared antigen specificity: many TCRs with shared cognate antigen are not similar in their sequences, and single mutations in TCR sequences can ablate antigen binding (see for instance Refs. [53–57] for early mutagenesis studies of TCR binding and Refs. [47, 48] for computational calculation of TCR sequence similarity within epitope-specific repertoires).

Clonal dynamics offer a complementary view to sequence similarity for the identification of T cell clones responding to a stimulus. Upon T cell expansion following antigen recognition, the frequencies of responding T cell clones in blood increase. As the TCR can be used as a barcode to identify single T cell clones, clones that expand and contract upon an immune stimulus can be identified and tracked over time through longitudinal TCR sequencing [13, 58–60]. TCR sequencing has proven successful in tracking the T cell response to a variety of stimuli, such as vaccines against infectious diseases [13, 61], cancer vaccines [12, 14], infection [8, 10, 62] and in evaluating repertoire fitness during chronic infections and recovery [63]. The identification of clones with shared dynamics can also be used as a first step in the identification of antigen-specific T cells, which can then be validated experimentally.

In the remainder of this article, we will focus on computational methods of identification of responding T cell clones through longitudinal TCR repertoire sequencing and the technical challenges that still need addressing.

## III. ACCOUNTING FOR NOISE IN LONGITUDINAL TCR SEQUENCING

### A. Estimating experimental and biological noise

One of the main challenges of analysing longitudinal TCR sequencing datasets is to separate changes in clonal frequencies due to a response to the stimulation of interest from background changes resulting from biological or experimental variability (Fig 1a, b). This task is made more difficult by the large number of clones in the repertoire, only a small subset of which are expected to respond to the challenge under study, against a background TCR repertoire that is largely stable over time [58–60]. However, even background clones are observed to change in a TCR sequencing experiment. Background variability can be classified into two main categories: (1) experimental noise, which consists of expression, sampling, and amplification noise [64–66]; and (2) biological variability corresponding to the population dynamics of clones [67, 68]. Experimental noise can be estimated from same [13, 66] or similar [69] timepoint replicates. However, repeat sequencing at each timepoint is often not available due to sample limitation and sequencing costs. Biological variability corresponds to the “neutral” dynamics of T cell clone frequencies when no known immunological signal is given over time scales of months to years [68].

**FIG. 1.**
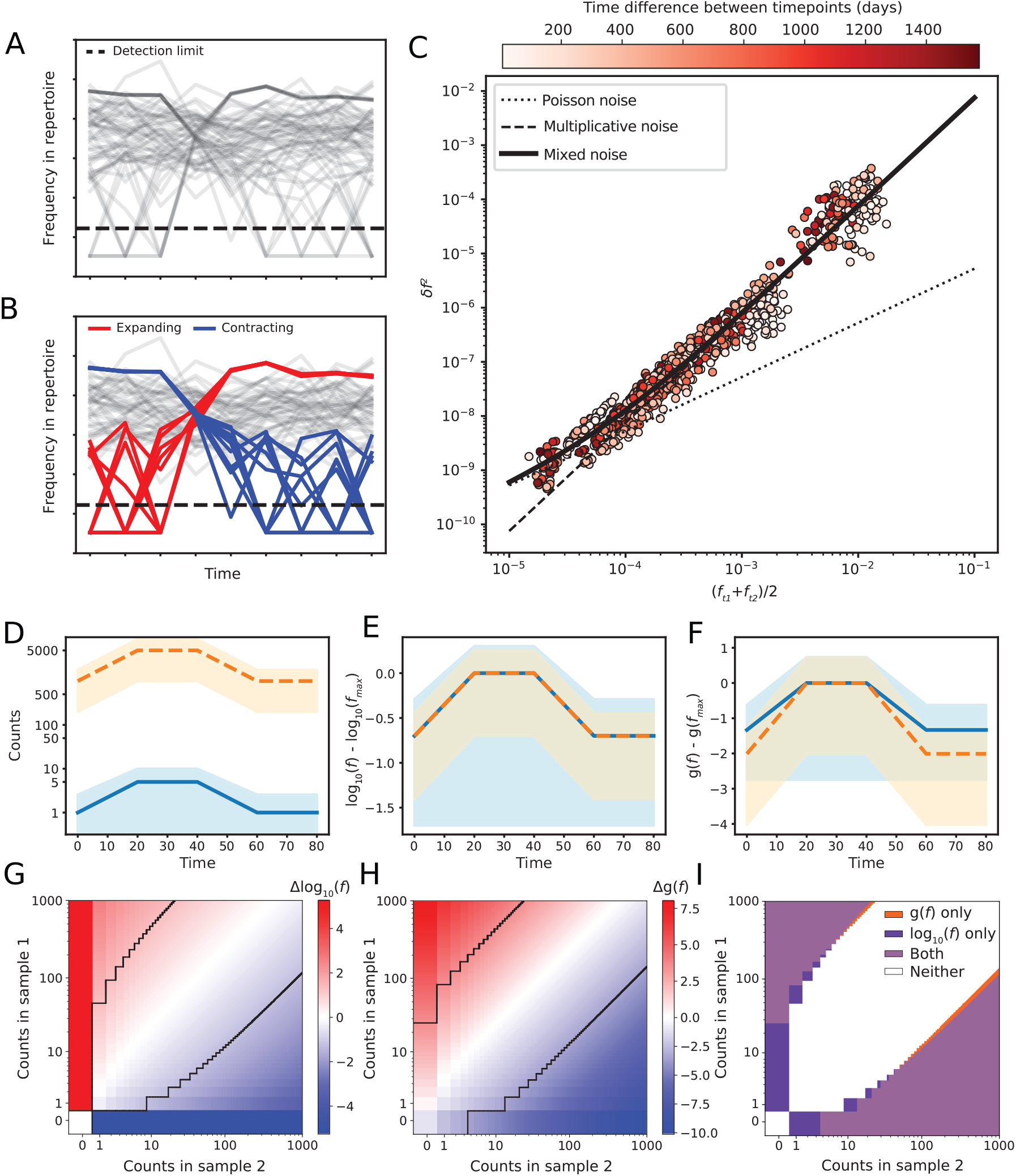
Noise-based transform of TCR frequencies. **A, B)** Schematics of the task on mock data: from a repertoire of unselected TCRs captured by longitudinal TCR sequencing (**A**), we want to identify TCR clones that are responding (red and blue) compared to background (grey) as in **B. C)** Strategy for calculating the *σ* and *β* parameters for *g*(*f* ). Fluctuations between timepoints (*δf* ^2^) are calculated for patient 11 from the PDAC-Vax cohort. Clones are binned by their mean frequency in the pair of timepoints, and the squared difference between the two frequencies in the pair is averaged in each bin and shown as *δf* ^2^. Multiplicative (*δf* ^2^ = *σ*^2^*f* ^2^, *σ* = 0.87) and Poisson (*δf* ^2^ = (*β/N*_*r*_ )*f, β/* ⟨N_*r*_⟩ = 5.3 *×*10^*−*5^) noise are shown as dashed and dotted straight lines. The fit of the mixed noise model of Eq. 2 is shown as a solid black line. **D-F)** Illustration of the effect of the noise-based transform on two simulated clones (orange and blue). The noise profile of each clone is calculated as 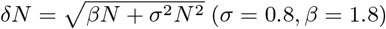. *δN* is shown by shading between *N* +*δN* and *N −δN*, with a minimum value for *N −δN* of 0. Trajectories of (**D**) raw counts, (**E**) log frequencies (with *N*_*r*_ = 5 *×* 10^4^) and (**F**) *g*(*f* ) (with *σ* = 0.8, *β* = 1.8, *N*_*r*_ = 5 *×* 10^4^). Both log_10_(*f* ) and *g*(*f* ) are normalised by subtracting the maximum. For the log-frequency plot, lower count bounds below 0.1 were replaced with a pseudocount of 0.1 before transformation for plotting purposes. **G-H)** Selection of expanding (red) and contracting (blue) clones between two timepoints (*N*_*r*,1_ = 7 *×* 10^4^, *N*_*r*,2_ = 3 *×* 10^4^) using (**G**) a fold-change threshold with |Δ log_10_(*f* )| *>* 1.3 or (**H**) |Δ*g*(*f* )| *>* 3.5 (*σ* = 0.8, *β* = 1.8). No pseudocounts are used for log_10_(*f* ) and Δ log_10_(*f* ). For comparisons involving counts 0, Δ log_10_(*f* ) is capped to twice the maximum absolute Δ log_10_(*f* ) calculated from all other comparisons. Thresholds are shown in black. **I)**: Overlap and differences between the two selection methods.

When analysing longitudinal TCR sequencing datasets, both of these effects compound, making the identification of true responding clones from background fluctuations challenging. Experimental noise is a particular concern for low-frequency clones, close to the minimal detectable frequency 1*/N*_*r*_ (where *N*_*r*_ is the sampling depth), which are strongly affected by sampling noise and for which estimation of frequency changes over time becomes challenging. By contrast, estimation of large clone frequencies can be done with high confidence, making the analysis of frequency changes relatively more reliable.

To illustrate how frequency fluctuations between sequenced samples depend on clone size in a real dataset, we analyse the properties of the fluctuations in the TCR frequencies of a patient (PDAC-Vax patient 11) with pancreatic ductal adenocarcinoma (PDAC) who received a personalised cancer vaccine as part of a Phase I clinical trial at Memorial Sloan Kettering Cancer Center [12, 14]. Patients in this trial had their tumour resected and were then treated with a single dose of atezolizumab (*α*PD-L1), followed by vaccination with 8 doses of a personalised vaccine containing antigens derived from the patient’s tumour, a standard 12-week cycle of chemotherapy and a single booster dose of the vaccine. We plot the mean squared difference in the frequency of the same TCR between any two timepoints (*δf* ^2^) as a function of its average frequency *f* in the two timepoints, averaged over all TCRs and all pairs of timepoints available for this patient (Fig 1c). TCRs are aggregated into frequency bins for each pair of timepoints based on every 10^th^ percentile of the average frequency distribution for that pair of timepoints. The observed *δf* ^2^ represents both experimental noise and biological variability, as both will impact the observed frequencies at each timepoint. The estimation is carried out on all clones that are detected in a minimum number of timepoints, to avoid over-inflating for small clones close to the detection threshold and reduce computational time. We find that at lower frequencies, *δf* ^2^ scales like the frequency *f* (dotted line), while at larger TCR frequencies, it scales like its square *f* ^2^ (dashed line). The first regime is consistent with sampling noise, where sequence count fluctuations follow a super-Poissonian law, *δN* ^2^*∼ βN*, where *β >* 1 corresponds to the amplification of sampling noise from single cells to the preparation of the sequencing library. The second regime, *δN* ^2^ *∼σ*^2^*N* ^2^, corresponds to multiplicative noise that may include experimental factors such as PCR amplification or true biological proliferation of the clone between the two timepoints. These two sources of variability propose the following form for sequence count variability:

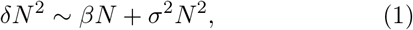

which can be written in terms of the frequency *f* = *N/N*_*r*_ (where we recall that *N*_*r*_ is the total number of reads) as:

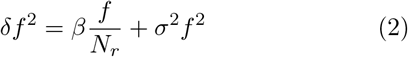

The *β* and *σ* parameters can be fit to observed fluctuations from the time series (Fig 1c, solid line) and are found to be *σ* = 0.87 and *β* = 2.6 for PDAC-Vax patient 11.

To understand the generalisability of this observed relationship, we extend the analysis of *δf* ^2^ to other 15 patients with PDAC who received personalised cancer vaccines (PDAC-Vax cohort, [12, 14]), as well as 3 other cohorts: a second vaccination cohort (YFV-Vax) and two natural infection cohorts (the Minervina-COVID and the COVIDsortium cohorts). The YFV-Vax cohort comprises 6 individuals who received a single dose of the live attenuated yellow-fever 17D vaccine [13] (YFV-Vax). A timepoint preceding vaccination is available for all individuals. The Minervina-COVID cohort includes 2 individuals who experienced mild SARS-CoV-2 infection and for whom longitudinal TCR sequencing was performed starting 15 days after infection [8]. A timepoint preceding infection is available for both individuals. The COVIDsortium cohort includes 46 healthcare workers who were exposed to COVID-19 in March 2020 in London [10]. Healthcare workers were tested weekly for COVID infection by PCR and date of first PCR+ is known. 6 individuals never became PCR+ and were used as healthy controls, while the remaining 40 experienced mild COVID-19 infection. For the YFV-Vax and Minervina-COVID datasets, two biological replicates are available for each sample. We consider each replicate as a separate time series and analyse two time trajectories for each individual. These cohorts were generated by different groups, using different sequencing technologies over a number of years and with different time intervals between samples and thus represent a range of different data types that the TCR community is faced with. By plotting the profile of background variability for each individual in each cohort (Figs S1 and S2), we find that the observed relationship between *δf* ^2^ and *f* is reproducible across individuals, cohorts, and sequencing platforms.

Sampling can be very different across cohorts and experiments, and proposed methods for background noise estimation need to be robust to lower sampling. We used the same data to illustrate how sensitive *β* and *σ* estimation is to the number of available samples and the sequencing depth (*N*_*r*_) of the available samples. This is an important consideration as deep or extensive TCR sequencing is not often available due to sample or cost limitations. We took the 17 timepoints for PDAC-Vax patient 11 and estimated the *β* and *σ* noise parameters for all combinations of 2 to 10 samples from the time series (Fig S3a, b). Overall, the median estimated parameters are similar and approaching the value estimated for the full time series (dashed red lines) for all sample subsets, but the estimation becomes less sensitive to sample selection (smaller spread) as more samples are included. To showcase the impact of sequencing depth (*N*_*r*_), we downsampled each sample in the time series to progressively smaller fractions (0.95 to 0.1) of its original sequencing depth (*N*_*r*_), 10 times for each fraction (Fig S3c, d). We find that *σ* monotonically increases towards its expected value from the full time series (red dashed line) with deeper sequencing (Fig S3c), while the estimated *β* is constant across all subsamples and in agreement with the expected value from the complete dataset (Fig S3d). Overall, we find that *β* and *σ* estimations are less impacted by sequencing depth than by number and selection of included samples.

Finally, we discuss how constant the estimated values of *σ* and *β* are within and across sequencing cohorts. These cohorts vary in sequencing depth, number of samples, sequencing platform, type of individual and immunological insult. We calculated the *σ* and *β* parameters for each individual time series by fitting the noise fluctuations (Figs S1 and S2). By looking across all individuals in each cohort, we find that the estimated *β* and *σ* parameters are similar within a cohort, and are of the same order of magnitude across cohorts (Fig S3e, f), suggesting that these parameters can capture both sequencing platform-specific technical variance as well as expected biological fluctuations. We find that *σ* has a median value of 0.58 and *β* has a median value of 1.91. Since *σ* and *β* estimations are similar within (and somewhat across) cohorts, these values can be used as general estimated noise parameters for TCR sequencing for any newly generated cohorts, to mitigate the impact of availability of sequencing samples or sequencing depth when lower sampling is available.

Overall, this analysis confirms that the background noise is larger than what we expect from sampling, and it includes other effects including multiplicative noise arising from biological and experimental variability.

### B. Detecting frequency changes between two timepoints

Responding clones are defined as clones whose frequencies change between timepoints in a manner that cannot be explained by background noise. The methods normally implemented for the identification of responding clones depend on whether replicate information is available for the cohort.

When a single sequencing sample is available for each timepoint, direct estimation of sampling and experimental noise is not possible. Instead, statistical tests can be used to establish whether a change in counts of a single clone is “surprising”, given the underlying sequencing depth. One simple way to do so and find significantly expanded or contracted clones is to use Fisher’s exact test. The Fisher exact test compares the difference of counts at two timepoints against their expected sampling fluctuations when no real change is present. To take into account the level of expected fluctuations, the test can be further corrected by directly editing the contingency table to hard-code a minimum fold-change difference. For instance, CloneTrack [12, 14] uses a corrected Fisher’s exact test with a fold-change cut-off of 2 and a strict p-value threshold to identify expanding TCRs with high specificity. To enforce the fold-change cut-off, the size of the control repertoire in the contingency table is divided by 2. A related method, also based on a Fisher’s exact test after count rescaling to account for technical noise, was implemented by Ayestaran and Blundell [69]. Alternatively, count differences observed between two samples can be directly compared to a Poisson null model of the noise to detect significant changes [10]. These classes of tests assume that the main source of variation is sampling noise, and do not account for multiplicative noise (*δf ∝ f* ).

When experimental replicates at the same timepoint are available, a more refined null model for background variations can be estimated from these replicates. Earlier methods developed for this case include adaptations of differential gene expression algorithms and Bayesian methods. Differential gene expression detection algorithms, such as edgeR [70], can be retooled to treat each TCR as a single gene. Timepoint replicates are used to fit a negative binomial form for the background noise, with an adjustable dependence of variance versus frequency. A significance test is then run to estimate whether the count difference between the two timepoints of interest is well above expected fluctuations [8, 13, 61]. Alternatively, custom Bayesian methods have been developed to specifically handle repertoire data. NoisET [65, 66] first learns a noise profile with variance *δf* ^2^ = *f/N*_*r*_ + (*σf* )^*β*^, where *β >* 1 is an adjustable exponent. It then computes for each TCR a posterior probability of response given an arbitrary prior on the number and typical fold change of responding clones. NoisET was applied to vaccination [13, 61] and infection [8] repertoire data, and was also adapted to infer the rate of change of clonal frequencies in T and B cell repertoires over long time-scales [68, 71].

Because most of these methods ignore biological variability, which, as we have discussed, induces multiplicative noise, a strict fold-change threshold is often used to enforce recognition of true frequency changes. Moreover, to avoid calling spurious differences in small clones close to the detection limit significant, small clones are often excluded from the analysis. These filtering steps increase the specificity of the methods (fewer false positives), but bias the identification towards large clones that undergo large frequency changes. More accurate knowledge of the underlying noise model allows us to build a better metric to represent frequency changes over time.

A common way to impose a fold-change threshold for selection of expanding clones is to use a log-fold change. As we will discuss, the log fold-change metric is actually the natural way to account for multiplicative noise. However, our previous discussion of the observed noise in TCR sequencing samples can be used to build a more principled transformation that can take into account the changing noise that TCRs at different clone sizes are subject to.

To compare differences between timepoints against a common threshold to define expansion across clone sizes, we want to build a function of the frequency *g*(*f* ) whose fluctuations remain constant as a function of *f* . This will allow us to compare differences in Δ*g* = *g*(*f*_2_) *−g*(*f*_1_) (where *f*_1_ and *f*_2_ are the frequencies at the two time-points) against a constant level of fluctuations which we set as significant. We can write the change in *g*(*f* ) upon variation of *f* as *δg∼ g*^*′*^(*f* )*δf* . In the multiplicative noise model, *δf ∼σf* . We obtain *δg∝ g*^*′*^(*f* ) *×σ× f*, which we want set to a constant. Thus, *g*^*′*^(*f* ) *∝* 1*/f* . A simple solution to this equation is a logarithmic transform, *g*(*f* ) = ln(*f* ). This leads to consider the log fold-change Δ*g* = *g*(*f*_2_) *− g*(*f*_1_) = ln(*f*_2_*/f*_1_) as an expansion score, which can be compared to a fixed threshold to call significant responding TCRs. This rationalizes the commonly used choice of log fold-change as a criterion for significant expansion.

However, as we have discussed, analysis of existing datasets suggests that, while multiplicative noise can explain the variance *δf* ^2^ observed at large clone sizes, smaller clones are dominated by Poissonian noise scaling, *δf* ^2^ *∝ f* . In addition, log-fold change calculation is challenging for clones that are not detected at the baseline timepoint, as an arbitrary value must be imputed for the missing measurement for the log-fold change to be calculated. The reasoning above can be extended to the more general form of the noise of Eq. 2, which leads to solving the equation *g*^*′*^(*f* ) = (*βf*/*N*_*r*_ + *σ*^2^*f* ^2^)^*−*1/2^, of solution:

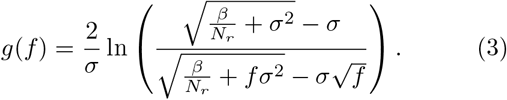

To align the transforms at high frequencies for all values of the sequencing depth *N*_*r*_, the integration constant is chosen to set *g*(*f* = 1) = 0.

We compare the theoretical behaviour of the log-fold change and the proposed *g*(*f* ) transformation for a large and a small clone and their fluctuations (Fig 1d-f). We find that in log-transformed space, frequency estimations for small clones have a much larger error compared to large clones (Fig 1e; fluctuations indicated by the shaded area for each clone), while the fluctuations in *g*(*f* ) space are comparable for the large and small clone (Fig 1f). Another desirable feature is that the *g*(*f* ) transformation gives more weight to changes in larger clones, consistent with the higher confidence we have in their fold change. We examine how the identification of expanding sequences between two samples with this transform compares with a log fold-change threshold. Consider two example repertoires *i* of total sequencing depth (*N*_*r,i*_), *N*_*r*,1_ = 70, 000 and *N*_*r*,2_ = 30, 000. We plot the log-fold change (Fig 1g) and Δ*g*(*f* ) scores (Fig 1h) for each clone count combination (*n*_1_, *n*_2_) from 0 to 1, 000. We also draw an equivalent selection threshold for the two in a black line: |Δ log_10_(*f* ) |*>* 1.3 or |Δ*g*(*f* ) |*>* 3.5. The selection thresholds were arbitrarily chosen to select similar regions of the count space. The two transforms show similar trends, identifying expanded (red) and contracting (blue) clones. To compare the count combinations that would pass the selection threshold for the two methods (Fig 1i), we identify regions of the count space that would be selected by none of the methods (white region), both of them (light purple region), only |Δ log_10_(*f* ) | *>* 1.3 (purple region), or only |Δ*g*(*f* ) | *>* 3.5 (orange region). Compared to fold-change, Δ*g*(*f* ) selects clones with a larger relative change at small clone sizes, and naturally places a count threshold for the expansion of previously undetected clones. By contrast, selecting based on log fold-change calls expanded all the clones that were not observed in the first timepoint.

In summary, while methods exist that can identify TCRs that vary across timepoints beyond the expected noise, they often only account for one noise regime. The proposed data-driven transform *g*(*f* ) allows for a single test accounting for both regimes of noise: sampling noise that dominates at low counts, and biological noise that dominates at large counts.

## IV. IDENTIFICATION OF RESPONDING CLONES FROM TIME SERIES

The methods reviewed in the previous section focus on the task of calling responding clones from a pair of timepoints previously identified as relevant, e.g. before and after vaccination or infection. These approaches are compatible with experiments where only two or few time-points are sequenced in a targeted way, and clones with specific dynamics need to be identified. What to do is less clear when there are more than two timepoints, and we have no prior knowledge on which timepoints to consider. In these cases, a more unbiased approach that does not make assumptions about the expected dynamics might be preferable. For example, unexpected clonal dynamics have been observed during COVID infection where, following a first wave of COVID-specific responders, a second wave of clones started expanding after 15 days [8, 62]. Such dynamics could only be detected thanks to two key differences compared to previous studies: (1) denser, and occasionally prospective, sampling was available and (2) a more unbiased approach was used for the detection of responding clones. As more and denser TCR sequencing datasets are published, an unbiased search for responses may unlock new insights into new T cell biology and drive important therapeutic advances. We review existing options for unbiased identification, and propose an improved approach.

For unbiased detection of responding TCR sequences, equivalent procedures as outlined above can be applied to all pairs of timepoints to look for any significant expansion or contraction at any point in the trajectory [10]. This method suffers from having to test multiple hypotheses (which grow quadratically with the number of timepoints), and thus decreasing statistical power, and may also miss features of the trajectories that cannot be summarized by pairwise comparisons, e.g. an initial peak at immunization followed by a rebound following a booster shot.

Instead of pairwise comparisons, Minervina et al. [61] proposed to project the trajectories of TCR clonal frequencies into a low-dimensional representation using principal component analysis (PCA), extending this idea to also include T cell phenotypic subsets (e.g. memory and naive). If and when clones clearly separate into distinct clusters, they can be classified into classes corresponding to different types of trajectories: steady, increasing, decreasing, or other more complex dynamics [8, 61, 62]. A clustering procedure was also performed without PCA, based on the distances between trajectories [72].

Unlike approaches based on pairs of timepoints, these methods do not directly account for noise, leading to biases and errors depending on which choices of thresholds and clone inclusion are retained. We suggest that the benefits of an unsupervised, unbiased search for changes can be combined with accurate accounting for biological and experimental noise by adequately transforming the clone frequencies and learning the expected variation in the background (Fig 2a).

**FIG. 2.**
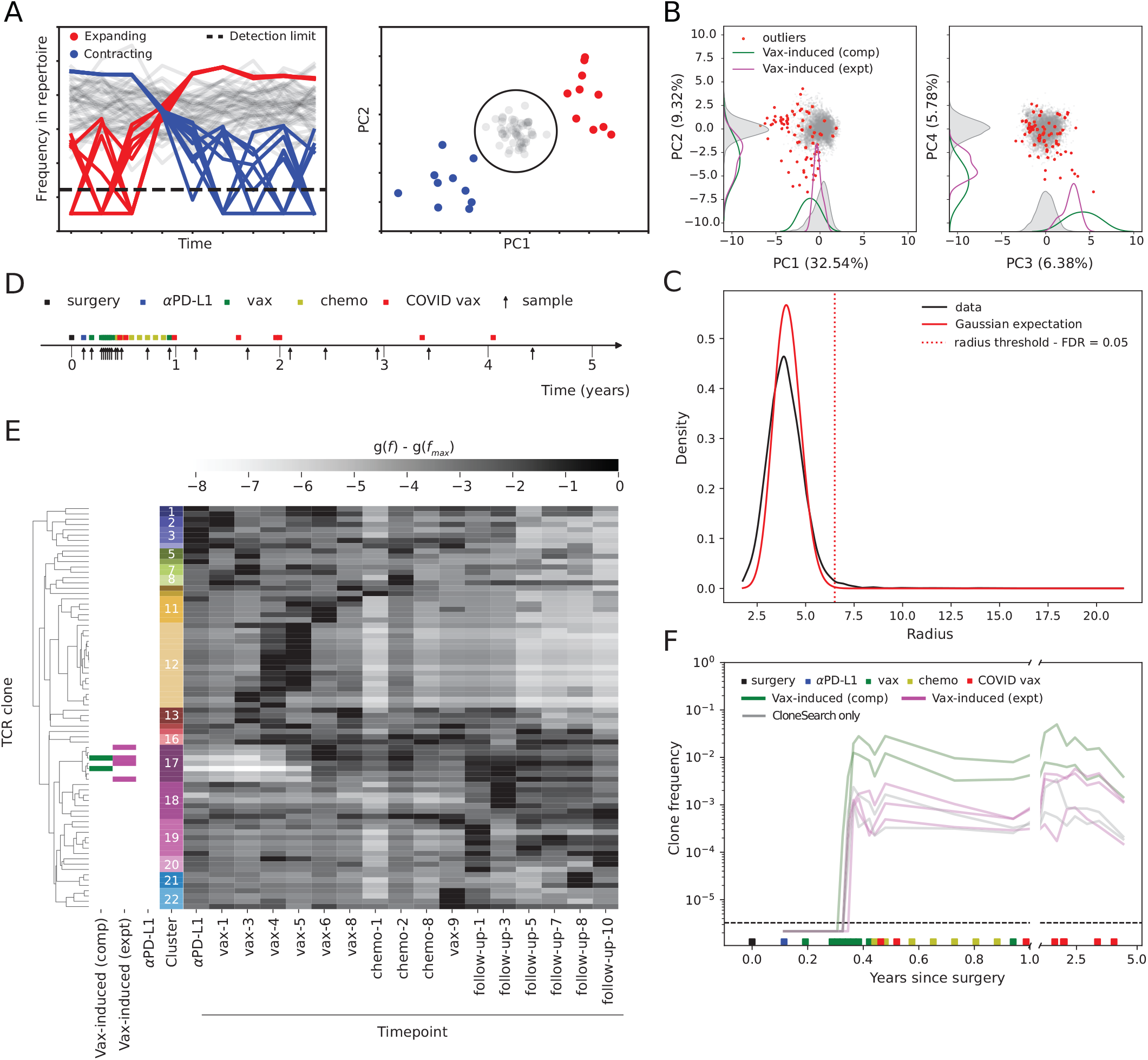
Identification of bulk and responding TCRs in PDAC-Vax patient 11 with CloneSearch. **A)** Illustration of CloneSearch outlier selection by Principal Component Analysis (PCA) followed by identification of outlier trajectories on simulated data, extending the example in Fig 1. B) Example PCA on PDAC-Vax patient 11. Only PCs 1-4 are shown. The density plots along the axes show the distribution of all (grey) and previously annotated clones: clones computationally identified by CloneTrack (Vax-induced comp, green) and experimentally identified (Vax-induced expt, pink). TCRs identified as outliers are indicated with red dots. **C)** Expected (red) and observed (black) distribution of radii from the centroid in normalised PCA space (*M* = 17 timepoints for this time series). The critical radius for outlier selection (dotted line) was chosen to achieve a false discovery rate of 5%. **D)** Treatment and sampling timeline for PDAC-Vax patient 11. **E)** Hierarchical clustering of identified outliers with annotations for previously identified sequences. Clusters are numbered and represented by different colours. Timepoints are ordered by time and follow the timeline in **D. F)** Trajectory plots for TCRs in cluster 17 from **E**.

Studies of healthy repertoires have shown that the TCR repertoire is mostly stable over time [58–60, 68]. Repertoire analyses of the T cell response to infection and vaccination have shown that only a small fraction of the repertoire responds to antigen stimulus [12–14, 61]. In fact, experimental and computational estimates suggest that, for any one pMHC, only 10^*−*6^ to 10^*−*4^ of T cells in the naive repertoire get activated and proliferate [39, 41, 73–78]. Thus, we expect the longitudinal time trajectories of T cell clones to be mostly stable over time, while a minority of responding T cell clones, which will expand upon antigen stimulation, are expected to show “unusual” trajectories.

Following Ref. [61], we reasoned that, if most TCRs are indeed constant over time, when each TCR trajectory is projected by PCA, we would see a dense bump representing background TCRs which are constant over time, and a few outliers corresponding to the responding clones (Fig 2a, b). To study whether this behaviour is in fact what observed in real data, we calculated the PCA projections of all clones that pass the clone detection requirement from PDAC-Vax patient 11, after transforming all frequencies with the *g*(*f* ) transform. To achieve this, we fitted the parameters *β* and *σ* from the time series of 53,243 TCRs across *M* = 17 timepoints in patient 11 as in Fig 1c. Using these parameters, we transformed each TCR frequency in each sample with the *g*(*f* ) transform of Eq. 3, and subtracted the maximum of each TCR over time, *G*(*f* ) = *g*(*f* ) *−g*_max_(*f* ), so that small and large clones are aligned in their maxima. We then projected the vectors (*G*(*f*_1_), *G*(*f*_2_), …, *G*(*f*_*M*_ )) of each TCR using PCA (Fig 2b, only the first four PCs shown; distribution for all components in Fig S4). Each principal component was centred and scaled to unit variance. Consistent with our hypothesis, in the space of the top principal components, we identify a dense sphere of points centred at the origin (in grey). This dense region represents the majority of the TCR repertoire, which has trajectories that are mostly constant over time. However, we also identify an additional cloud of outliers. The presence of these outlier TCRs which have non-constant trajectories is consistent with the fact that this patient received *α*PD-L1 treatment, cancer vaccination and COVID vaccines during the time under observation. Thus, we expect the clones with outlier trajectories to be the T cell clones responding to any one of these stimuli. For patient 11, we have previously annotated clones known to respond to vaccination and identified either experimentally through peptide stimulation experiments (N = 6), or computationally by CloneTrack (N = 2) [14]. CloneTrack had also been used to search for clones responding to *α*PD-L1 treatment by comparing the relevant timepoints, but no TCRs expanding upon this treatment were identified for this patient [12]. Thus, we can check if the identified clones are among the clones with outlier trajectories. As expected, we find that previously identified clones (experimentally in pink, and computationally by CloneTrack in green) sit outside of the main density in multiple principal components (Fig 2b and Fig S4).

Since the unique trajectories of responding TCR sequences sit outside of the main distribution, we can use the main distribution to define the space of “expected” trajectories, and draw a threshold to identify the outliers. This threshold will be specific for the time series and depend on the number of samples as well as the typical distribution of the background in the time series. The background implicitly represents the noise of the sample, without a noise model being explicitly calculated. Because each principal component is a linear combination of transformed frequencies at multiple timepoints, we expect the spread across non-responding TCRs to be approximately Gaussian around the PCA centroid (the origin) according to the central limit theorem. To verify this, we measure the distance, or radius *r* of each TCR to the origin of the normalised PCA space. If TCRs were normally distributed around the origin, which we assume to be true for non-responding TCRs, we would expect the radial distribution *P* (*r*) to be given by:

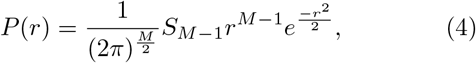

where *S*_*M−*1_ = 2*π*^*M*/2^*/*Γ(*M*/2) is the area of a unit sphere in *M* dimensions. We can then define a radius threshold above which we do not expect to observe any non-responding TCR under Eq. 4 as a null hypothesis, with a fixed false discovery rate (FDR) to account for multiple testing (see Methods, Fig 2c). This method allows us to identify responding TCRs as exceeding that critical radius, regardless of the timing and knowledge of the event which caused the response. The definition of the radius can be adjusted to be more or less stringent by tuning the FDR.

The 76 responding TCRs (FDR = 0.05) identified for PDAC-Vax patient 11 are shown as red points in Fig 2b and their trajectories are shown in Fig 2e, f and Fig S6. To help interpretation of the identified TCRs, responding sequences were grouped into 22 clusters based on their transformed trajectories (Fig 2e, Fig S6 and Methods), and compared to previously identified sequences for this patient (experimentally identified in pink, computationally identified in green; annotations for previously annotated sequences in Fig 2e on the left-hand side of the heatmap). By plotting the time trajectories of all TCRs in each cluster together with available clinical information for the patient (Fig S6) we are able to make hypotheses on the function of the identified responding clones. One of the main identified clusters corresponds to vaccine-induced clones (cluster 17): these clones are undetectable at the beginning of the time series, expand upon personalised cancer vaccination (green squares) and remain high up at the follow-up timepoints (Fig 2f), as previously described [12, 14]. In this cluster, we observe two out of two clones that had been previously identified by CloneTrack (indicated in green). Thus, by using an unbiased method which simply looks for “unexpected” trajectories, we can identify the complete set of clones that were previously found computationally in this patient. We also identify 5 other clones in cluster 17 with similar time trajectories to the CloneTrack-identified ones, albeit at smaller magnitudes. Of these, 3 had been experimentally detected, but not identified by CloneTrack. The lower magnitude of these experimentally identified TCRs is likely the reason they were missed by CloneTrack in the first instance. Finally, the method identifies additional trajectories of interest in the other clusters, the functionality of which remains to be mapped. Importantly, all of these time trajectories are identified in a single analysis, without specifically looking for a response at specific times and to specific stimuli.

The outlined approach addresses the need for a noise-aware transformation and the need for unbiased identification of outlier trajectories. We packaged all the steps required for this in a method that we call CloneSearch.

## V. UNBIASED SEARCH FOR RESPONDING T CELL CLONES

### A. Identification of known and new T cell responses

To showcase the power of an unbiased search for responding T cell clones, we re-analysed the four published datasets introduced above to answer two key questions: (1) can an unbiased method truly recapitulate previously-published results and (2) can it find previously-uncharacterised responses? For each cohort, we compared the overlap between previously captured and responding clones as identified by CloneSearch (Fig 3a-d). For the PDAC-Vax cohort (Fig 3a), three categories of previously identified clones are available: an experimental set of cancer vaccine-specific clones (“Vax-induced (expt)”) and two sets of computationally identified clones. These computational clones were found by designing two different statistical tests to specifically detect clones expanding during personalised cancer vaccine treatment (“Vax-induced (comp)”) and clones expanding during *α*PD-L1 treatment. We show the number of clones that are found (solid bars) or missed (hatched bars) and the percentage of previously identified clones found by CloneSearch for each category separately (Fig 3a and e, respectively) after a single run of CloneSearch, with no tailoring for detection at specific timepoints. Clones that had previously been identified computationally as responding to the vaccine are most easily recapitulated, while experimentally identified and *α*PD-L1-induced clones are more easily missed by CloneSearch (Fig 3a, e), likely because they expand less and have thus less characteristic trajectories.

**FIG. 3.**
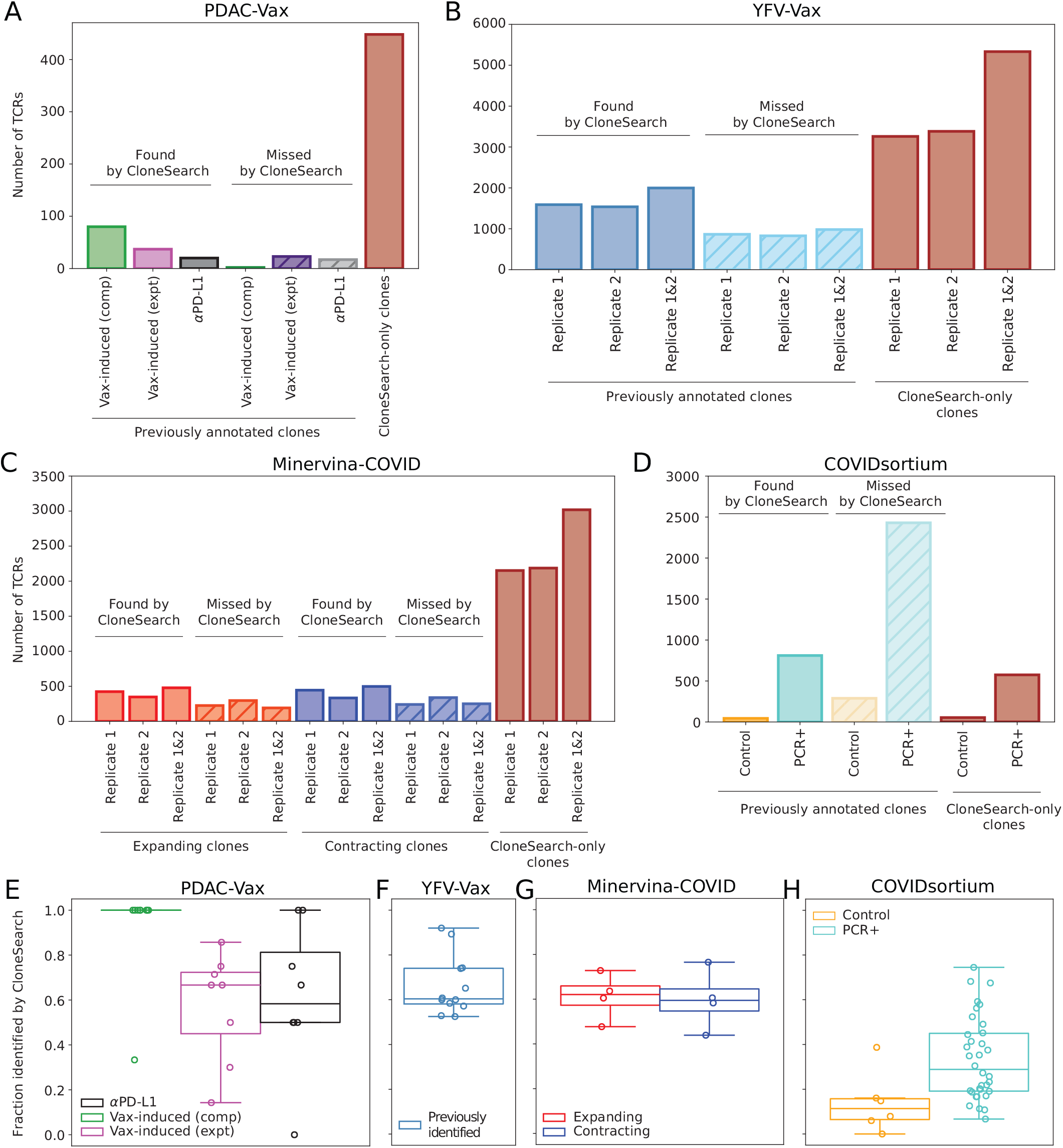
Overlap of identified outliers by CloneSearch with previously annotated clones across datasets. **A-D)** Overlap of CloneSearch-identified and previously annotated sequences in (**A**) PDAC-Vax, (**B**) YFV-Vax, (**C**) Minervina-COVID, and (**D**) COVIDsortium datasets. In YFV-Vax and Minervina-COVID, the two replicates available for each individual are analysed and reported separately (Replicate 1 and Replicate 2). Since previously identified clones were found using both replicates at the same time, we also show the overlap by merging the outliers from either replicate (“Replicate 1&2”). In the COVIDsortium dataset, 6 healthcare workers never became PCR+ for COVID infection. We thus split the cohort in Control (never PCR+) and PCR+ individuals. **E-H)** Proportion of previously identified sequences that are identified by CloneSearch, for each cohort: (**E**) PDAC-Vax, (**F**) YFV-Vax, (**G**) Minervina-COVID, and (**H**) COVIDsortium.

We repeat the overlap and recall analyses for previously identified clones in the 3 remaining cohorts. In the YFV-Vax cohort responding clones were originally defined as significantly expanding between days 0 and 15 post vaccination using an explicit noise model based on the available sequencing replicates (Fig 3b, f). In the Minervina-COVID cohort (Fig 3c, g), a PCA-based method on the largest clones at each timepoint had identified clones that expanded between days 15 and 37 postinfection (in red), and others that contracted between days 15 and 30 (in blue). Finally, in the COVIDsortium cohort, putative COVID-responding clones were identified in individuals who became PCR+ (teal) for COVID or remained healthy (orange) during the study by comparing all pairs of timepoints (Fig 3d, h). Overall, CloneSearch can easily identify vaccine-induced clones (Fig 3e, f), which tend to have more characteristic trajectories, while it is less likely to pick up infection-or *α*PD-L1-induced clones (Fig 3e, g, h), which tend to have less characteristic trajectory shapes. In all cases, Clone-Search finds *>* 50% of previously identified clones in most individuals (Fig 3e-h), with the exception of the COVIDsortium dataset.

The study of each cohort used different tests and significance thresholds for calling responding clones, resulting in varying rates of false positives. This might explain the different rate of clones missed by CloneSearch in each cohort (Fig 3e-h). We specifically looked at the missed clones from the COVIDsortium cohort, where CloneSearch had the lowest recall. In control individuals, who did not get COVID and should not experience clonal expansion, the original study still reported responding clones, which can be considered as false positives. Most of them are not called by CloneSearch, and CloneSearch generally does not identify many responding clones in controls relative to PCR+ individuals (Fig 3d, h), suggesting that CloneSearch de-noises the search for responding clones.

To further study the effect of FDR on CloneSearch outlier selection and recall of published sequences, we calculated the percentage of previously annotated sequences that are observed by using a range of radius thresholds corresponding to different FDRs (Fig S5, left panels). As expected, across the board, fewer previously annotated sequences are identified at more stringent FDRs. Interestingly, in the PDAC-Vax dataset (which used very stringent expansion definitions) the decay of the recall curve is slower for the computationally identified vaccine specific clones. Moreover, in line with our previous hypothesis that CloneSearch denoises the search for the COVIDsortium data, we find that the Control individuals in the COVIDsortium cohort have a steeper dropoff of identified previously annotated clones compared to the individuals who became PCR+.

We found that CloneSearch identifies many clones that had not been previously annotated in the published cohorts (Fig 3a-d, “CloneSearch-only clones”). The proportion of new clones identified at FDR = 0.05 is different across cohort (Fig S5, right panels), highest in the PDAC-Vax and lowest in the COVIDsortium cohort. These results are consistent with the way responding clones were originally identified in each of these cohorts. For the PDAC-Vax cohort, a very targeted search for clones responding to personalised cancer vaccines or immunotherapy was conducted with very strict thresholds, while for the COVIDsortium cohort all timepoints were evaluated in pairwise comparisons. Unbiased methods, like CloneSearch, are not limited to responses in specific timelines, but can pick up responses across the time series, which makes them less sensitive to choices made in the experimental protocols. Patients in the PDAC-Vax cohort were followed up for up to 6 years, and during this time they may have been subjected to other immunological insults that explain these newly identified clones. To showcase the power of an unbiased search for T cell responses, we showcase two case studies of newly-identified clones in two patients from the PDAC-Vax cohort (Fig 4). First, we look at the outliers identified for patient 11 from the PDAC vaccination cohort (Fig 4a). As described above, we find that the previously identified vaccine-specific T cell response (marked in green and pink) is contained (and expanded) within cluster 17. There is, however, another cluster, cluster 12, which contains T cells which are mostly undetectable at prevaccination timepoints and expand at vaccination doses 4 and 5 (Fig 4a, b). We use the available paired single-cell RNA sequencing dataset for this patient to assign CD4 and CD8 phenotypes to the identified CloneSearch outliers. We find that cluster 17 is composed of CD8 T cells, as previously described [12, 14], while clones in cluster 12 are predominantly CD4 T cells. These findings suggest that personalised mRNA vaccination stimulated both a CD8 and a CD4 response in this patient, although the CD4 response could not be picked up by previously used methods, likely due to its smaller magnitude.

**FIG. 4.**
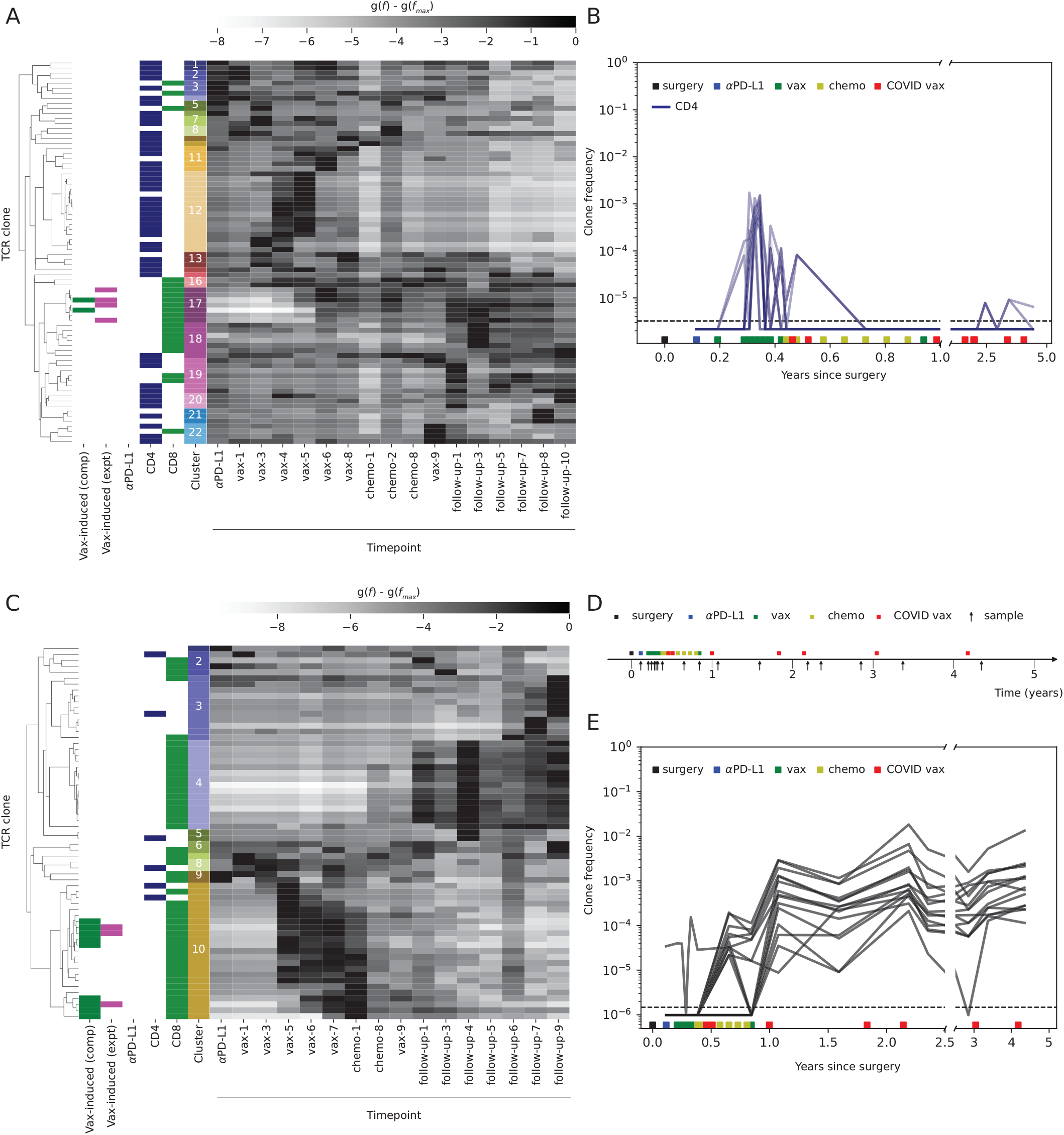
Identification of putative CD4 and COVID vaccine responses in the PDAC-Vax cohort. **A)** Heatmap of 76 CloneSearch outliers for PDAC-Vax patient 11 (as in Fig 2) annotated with CD4 and CD8 phenotype from matched single-cell sequencing. Cluster 12 contains CD4 T cells that seem to peak early during the vaccination timeline. **B)** Trajectories of CD4 T cells from patient 11 which are the putative CD4 vaccine-induced response from cluster 12 in **A. C)** CloneSearch outliers (*N* = 63) identified in PDAC-Vax patient 10. Like for patient 11 (**A** and Fig 2), clones that had been previously characterised as induced by vaccination or by *α*PD-L1 treatment are annotated in the heatmap. Cluster 4 contains CD8 T cells that seem to respond to COVID vaccination. **D)** Treatment and sampling timeline for PDAC-Vax patient 10. **E)** Trajectories of T cells from patient 10 that are the putative COVID vaccine response (cluster 4). COVID vaccination timepoints are indicated by the red squares on the x-axis.

We then looked at the outliers identified for PDAC-Vax patient 10 (Fig 4c). As in patient 11, previously identified vaccine-specific responses are contained within a single cluster, cluster 10. We also noticed a large cluster of T cell clones (cluster 4) with a characteristic shape (Fig 4e). By mapping the available clinical information for this patient on their sampling timeline (Fig 4d), we find that clonal expansion in this cluster roughly corresponds to timepoints of COVID vaccination (indicated by the red squares in Fig 4d). To validate the hypothesis that these clones are COVID-specific, we tested whether cluster 4 contains sequences that are more similar to known COVID-reactive sequences than background sequences, but could not detect any enrichment (Fig S7). COVID reactivity for these TCRs thus remains to be validated, and targeted experimental approaches would be needed to address this question. Alternative hypotheses can also be considered to explain the observed behaviour of these TCRs: vaccination or respiratory infection not reported in the clinical data, or epitope spread from the activity of the immune system against the tumour [79, 80]. Thus, the ability of unbiased methods like CloneSearch to identify TCRs responding to immunological events that are unexpected opens avenues for generating hypotheses and identifying novel responses that can be validated experimentally.

### B. Comparison of log transform and data driven transform of frequencies in outlier selection

We have so far discussed two separate improvements to the identification of TCRs that clonally expand in response to stimulus. The outlier identification strategy, however, can be applied as easily with either transformation. We thus compared our proposed noise-based transform *g*(*f* ) and the classically implemented log transformation of frequencies in the PCA projections and outlier identification, to establish whether a better noise transformation improved recall of expected responding TCRs. We can do so by comparing CloneSearch-identified TCRs by either method with previously annotated TCRs in each cohort. A direct overlap measure between the identified outliers and the published sequences is sensitive to threshold selection and, as we have discussed, the different published datasets have different false discovery rates. Thus, we perform this comparison by labelling the TCRs in the dataset as previously annotated (or not) and ranking them by their radius and calculate the area under the receiver-operator characteristic curve (AUROC) to establish whether previously annotated sequences were more likely to be classified as outliers using either the *g*(*f* ) or the log-transform, treating previous annotations as “ground truth” (Figure 5a-d).

**FIG. 5.**
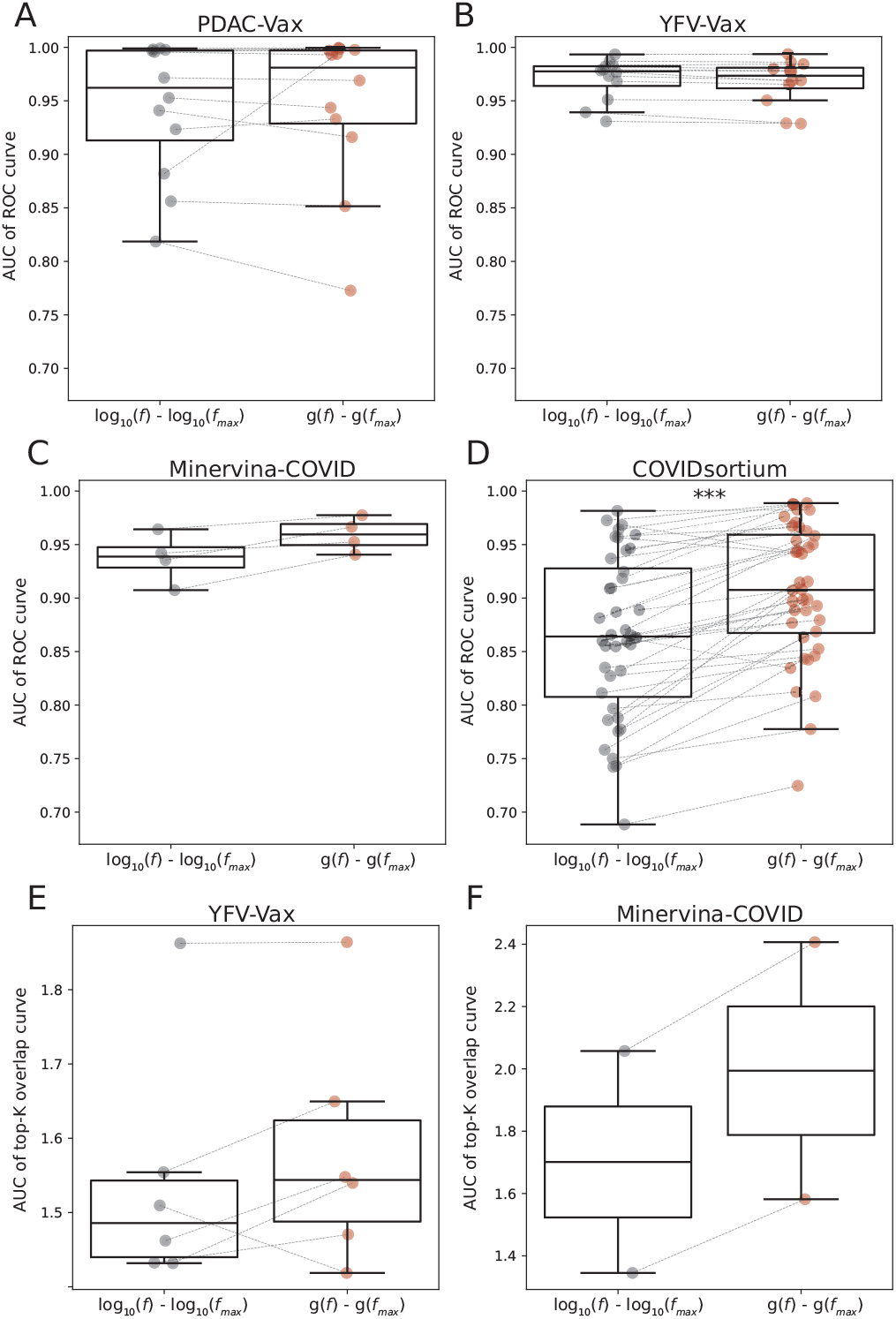
Comparison between *g*(*f* ) transform and log-transform. **A-D)** Area under the curve (AUC) of the receiver-operator curve (ROC) generated by ranking all TCRs by PCA radius *r*, as obtained using each transform, and la-belling with the previously published sequences as ground truth for each dataset: (**A**) PDAC-Vax, (**B**) YFV-Vax, (**C**) Minervina-COVID and (**D**) COVIDsortium. **E-F)** AUC of the overlap index calculated between the top *K* (where *K ∈ {*1, 2, …, 1000*}*) clones ranked by radius *r* in the two replicates versus log_10_(*K*) (shown in Fig S8), for (**E**) YFV-Vax and (**F**) Minervina-COVID. This AUC serves as a measure of consistency between different sequencing runs, and thus of the robustness of the method. Stars indicate p-values calculated using the Wilcoxon paired test, with alternative hypothesis that the AUC from the *g*(*f* ) transform is greater than the AUC from the log-transform: *** *≤* 0.001 *≤* ** *≤* 0.01 *≤* * *<* 0.05.

In all cohorts except YFV-Vax, we find that the *g*(*f* ) transform increases the median AUROC relative to the log-transform. This difference is significant only in the COVIDsortium dataset (Fig 5d), which is both the hardest to recapitulate and has the most comparison points. We also note that, based on these four datasets, identification of vaccine-induced responses (Fig 5a, b) is easier than identification of infection-induced responses, which have lower AUROC in all models (Fig 5c, d). As discussed below, these results may be confounded by the number of samples in the time series for each individual in these cohorts, as well as the average sequencing depth of the samples.

For two cohorts, YFV-Vax and Minervina-COVID, we have access to two replicates at each timepoint for each donor, which we analyse as two separate time series. We can compare outliers across these two replicates to assess the robustness of the method to sequencing noise. To do this, we calculate the proportion of overlapping sequences in the top-K clones in each replicate, where K is varied from 1 to 1000 clones ranked by their outlier score *r* (Fig S8). We compute the area under the curve of this quantity as a function of log_10_(*K*)(Fig 5e and f, for YFV-Vax and Minervina-COVID, respectively, and Fig S8 for individual curves). We find that in both cohorts, the *g*(*f* ) transform leads to better consistency than the log-transform across replicates.

### C. Impact of sample selection and sequencing depth on identified responses

The number of samples that can be collected, the time interval between them, and the relevance of the sequenced timepoints to the underlying immunological events are often limitations to the detection of T cell responses from longitudinal TCR sequencing. The PDAC-Vax cohort offers an ideal setting to ask how the sample number and selection affect detection of T cell responses: for patient 11, 17 longitudinal TCR sequencing samples were acquired over the span of 5 years and events that may have caused T cell responses are known. Overall, we find that, when the task is to identify cancer vaccine-induced responses, as few as 2 or 3 timepoints framing the relevant vaccination timeline are enough for detection by outlier identification (see Appendix A 1 for full discussion).

Another concern in the identification of responding TCRs is sequencing depth: when is a sample too shallow for responses to be reliably detected? As expected, we find that outliers can be identified also at lower sequencing depths, although the number of identified responding TCRs is positively correlated with higher sequencing depth (see Appendix A 1 for full discussion).

Importantly, both sample selection and sequencing depth impact the TCR clones that are available for analysis, as smaller clones and clones that are detected at fewer timepoints are more likely to be missed when sparser or less deep sequencing is available. Throughout the presented analyses, we have used a clone filtering step to reduce spurious identification of small clone fluctuations as outliers. This is a common strategy in analysing TCR sequencing data: to avoid detection of noise, only clones that are either largest in a sample, or clones that have a minimum count are included. For instance, in Ref. [8], only clones that are in the top 1,000 most expanded clones in any one sample are included, and Refs. [12, 14] retain clones that are found at a minimum count of 3 in at least 2 timepoints. Throughout the analyses presented, we have used the latter clone filtering strategy, with a couple of exceptions (full details in Sections VII B and VII F). However, the filtering strategy is arbitrary, and the filtering step used for most of our analyses will remove clones that are large at one timepoint, but not detected in other timepoints in the time series. This strategy is not optimal when dense sampling is not available, as a clone detected at a single relevant timepoint might still carry a lot of information. We assess the impact of different clone filtering strategies and suggest that Clone-Search may provide a useful tool to establish the optimal clone inclusion criteria for each longitudinal sequencing experiment (see Appendix A 2 for full discussion).

### D. TCR characteristics of identified responding T cell clones in all cohorts

Lastly, previous work has explored whether TCRs responding to immune challenges have a high probability to be generated and found in naive repertoires [81–84]. To test whether the identified responding TCRs were distinct from the background, we compared the identified outliers to each donor’s own background (Fig 6). First, we calculated the length distribution of the CDR3*β* of identified outliers, compared to a size-matched background for each patient, finding no difference between the two (Fig 6a). We also estimated the probability of each TCR to be observed in a functional repertoire *P*_post_ [85– 87] (Fig 6b). We find that these parameters are comparable across identified TCRs and across cohorts. These findings suggest that, when one is agnostic to the antigen, no simple generic feature distinguishes responding TCR from background ones.

**FIG. 6.**
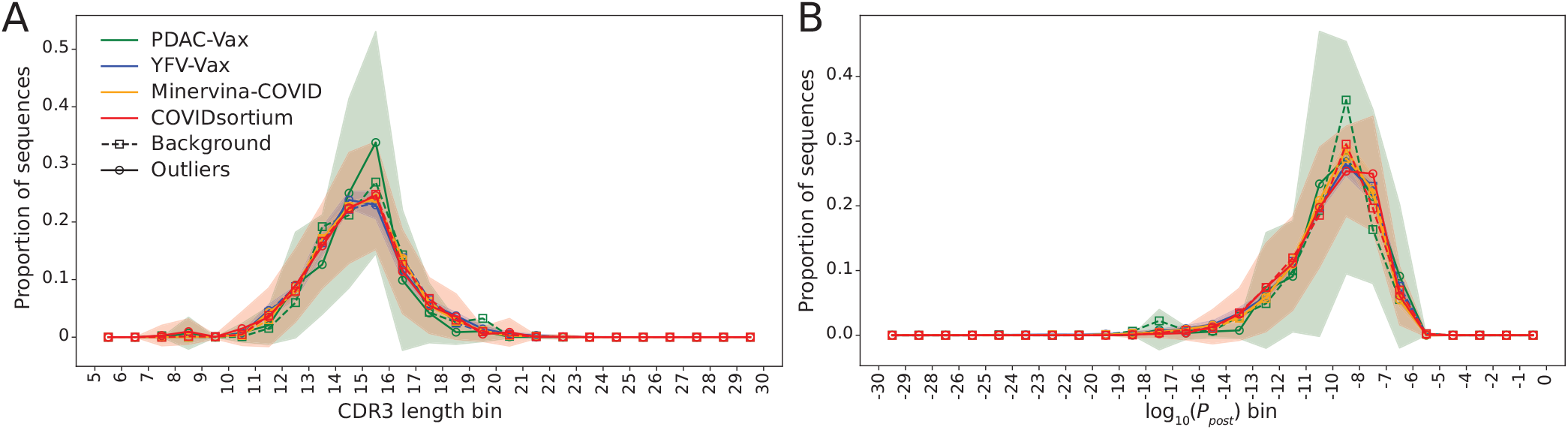
TCRs identified by CloneSearch are not different from background or between cohorts. (**A**) CDR3 length and (**B**) probability of selection (*P*_post_) calculated for the identified outliers and size-matched controls for all patients in each cohort. The average across patients in a cohort was calculated for outliers (solid lines, round markers) and size-matched background (dashed lines, square markers). Distributions are shown as histograms and each point on the plot represents the midpoint of the bin.

## VI. DISCUSSION

Longitudinal TCR sequencing provides a rich data source to profile the magnitude and duration of T cell responses to different stimuli. Compared to experimental measurements of antigen specific T cells over time, longitudinal TCR sequencing has a few advantages: (1) it does not require knowledge of the specific pMHC inducing the response, needed for *in vitro* T cell stimulation assays, (2) it can measure both magnitude and breadth of response and (3) can provide novel candidate TCRs for validation experiments. However, computational discovery of T cell responses suffers from one main challenge: the need to account for biological and technical noise. High-magnitude responses, such as those observed after vaccination, are more easily detected by traditional methods. However, smaller, subtler responses are difficult to disentangle from background fluctuations. Here, we have reviewed current methods that account for noise in TCR sequencing experiments, and proposed a novel way to account for noise directly in the transformation of TCR frequencies. The introduced *g*(*f* ) transform is parametrised in a time series-specific way to take into account the noise typical of small (Poisson scaling) and larger clones (multiplicative noise) and thus calculate a more meaningful expansion score (Δ*g*) between samples. Using this transform, we can identify TCR clones that expand or contract with confidence at both large and small frequencies. The noise profile is estimated directly from the sequencing datasets without the need for biological or technical replicates, in contrast with previously published methods [66, 88]. Parameter estimation for the noise profile can be carried out with as few as 2 longitudinal samples and is robust to sequencing depth, although performance stabilizes as more and deeper samples are included in the time series. We find that the inferred noise parameters, *β* and *σ*, are relatively stable between patients in a single cohort, and between cohorts (Fig S3), suggesting that a user-input value could be used for cohorts where samples are limited and parameter estimation challenging.

Our review of methods for identification of responding TCR sequences also highlighted that most methods only search for TCR expansion between specific timepoints. This approach is justified when few TCR sequencing samples are collected for each time series, and the timepoints are specifically selected to capture expected clonal dynamics. With the advent of larger sequencing cohorts, where donors are monitored over long time periods and with multiple sequential timepoints, more unbiased methods may prove more powerful. We have highlighted that unbiased clone identification can uncover new biology, and the development of methods that can robustly identify responding TCRs from a time series without a starting hypothesis is an important area of development. Therefore, we have paired the noise-aware transform with the identification of clones that have “unusual” trajectories compared to the background. The idea of clustering TCR trajectories to find distinct characteristic trajectory shapes had previously been proposed [8, 61, 62, 72]. Our proposed approach differs from previous approaches in two ways: first, these methods were only applied to the largest clonotypes in the repertoire, bypassing the need for a transformation that takes into account the different noise profile for smaller clones. Including smaller clones makes it possible to identify lower-magnitude responses. Second, we extend the PCA analysis to the concept of outlier identification, thus removing the need to identify a cluster similarity threshold and define the number of clusters of co-varying trajectories in the data *a priori*. We simply propose to identify all responding trajectories which are different from the background. The user can then assign their function based on their trajectories. Thus, we are also able to pick up smaller responses driven by only a handful of clones which may not form a well defined cluster. This hypothesis-generating approach is packaged in a new tool, which we name CloneSearch. The tool is made available for the community as a Python package at https://github.com/mm523/CloneSearch.

We demonstrated that CloneSearch can replicate results from previously published methods, but can also identify responses that had not previously been reported. CloneSearch-identified outliers did not look different in their sequence properties from background sequences, consistent with previous reports on antigenspecific repertoires [81]. Importantly, CloneSearch could identify responses to multiple stimuli within a single analysis, without any previous knowledge of the timing or shape of the identified response: the detection does not rely on the user knowing when to expect an observable response. Of note, CloneSearch could identify a CD4 response in PDAC-Vax patient 11, distinct from the CD8 response identified in the original study. This is remarkable as the statistical test previously applied to the dataset missed these clones as these are relatively low fluctuations. Interestingly, the dynamics of these clones are similar to what was recently described in primary COVID infection, where CD4 T cell clones are shown to expand in a distinct wave from CD8 clones [89]. In addition, CloneSearch highlighted a putative response to COVID vaccination in PDAC-Vax patient 10. Finally, CloneSearch can also be used with a more liberal threshold as a pre-screening method to select a set of clones of interest, which can then be further screened with methods with higher specificity to get more confident hits. This reduces multiple-hypothesis testing and may reduce the false negative rate.

Our analysis also highlights that sample selection is, and always will be, the major bottleneck to discovery of T cell responses. Despite the agnostic behaviour of CloneSearch to the time of stimulus, our analysis shows that the only clones that can be detected are the ones that are stimulated by insults which “scar” the repertoire around the time of sampling. Thus, not all combinations of samples from a time series will give equal performance. Moreover, computational identification of T cell responses is limited to responses that trigger a frequency change for a clonotype in a repertoire. Although re-activation of large clones may not lead to their expansion and thus will not be detected, recall responses to booster vaccination were shown to trigger a detectable expansion [61].

Even when no reported immune challenge affects the repertoire, T and B cell repertoires change over time. This evolution is slow. Clones have been shown to be stable over years or even decades [58, 59, 68, 71, 90]. These changes induce some variability, which is included in our noise model over the duration of the experiment. Beyond these slow fluctuations, we expect that, when longitudinal sampling is performed over long times, individuals may experience unreported or undetected immune challenges in addition to known events causing the expansion of some clones, as was shown in Ref. [69] in three healthy donors. Understanding the frequency and magnitude of these events in healthy donors remains an open question. By using unbiased methods, these responses can be identified. Indeed, we speculate that some of the outliers and their clusters that CloneSearch detected in these cohorts but to which we were unable to assign a function correspond to these unreported challenges and may not all be considered as false positives.

Overall, despite the many technical challenges, longitudinal TCR sequencing provides a powerful tool to quantify magnitude and duration of T cell responses to immunological events. CloneSearch provides a robust framework to identify responding TCR clones from longitudinal TCR sequencing datasets in a manner that is noise-aware and unbiased to the time of stimulus.

## VII. METHODS

### A. Datasets

The datasets analysed were all previously published. The pre-processed data from the Adaptive Biotechnologies and 10X single-cell sequencing pipeline for the PDAC-Vax cohort [12, 14] was obtained from the authors. The previously published data was further extended to capture more longitudinal timepoints with TCR Vb sequencing and more scRNAseq samples were collected for deeper phenotyping of the clonotypes of interest (paper under revision). The YFV-Vax cohort [13] is available from the NCBI Sequence Read Archive (SRA, accession no. PRJNA493983). The Minervina-COVID cohort [8] is also available from the NCBI SRA (accession no. PRJNA633317). The data preprocessed with MiXCR [91] for both these datasets was obtained from the authors. The raw data for the COVIDsortium cohort [10] is available from the NCBI SRA (accession no. PRJNA718557) and the data preprocessed by Decombinator [46, 92, 93] was downloaded from https://github.com/innate2adaptive/CovidsortiumTCRExpanded/. We retained only individuals for which at least 3 timepoints were available (40 of 46). For all analyses, the clones were identified using their TRBV, TRBJ and CDR3*β* nucleotide sequence. TCR frequencies were calculated using either the provided counts (PDAC-Vax) or UMI-corrected counts (YFV-Vax, Minervina-COVID and COVIDsortium), according to what provided by the preprocessing software.

### B. Sequence data processing

Each individual timeline was processed separately. As sample quality can affect frequency calculation and outlier detection, we removed samples with a total TCR count *<* 5000. For samples sequenced by Adaptive Biotechnologies (PDAC-Vax cohort), we also removed any samples that have *≤*15 productive sequences / ng of DNA to remove samples with poor sequencing quality.

We also applied a clone filtering step to remove small clones from the analyses. For consistency across donors and cohorts, the clone filtering is performed in the same way for all analysed time series. Unless explicitly stated otherwise, a TCR is kept for downstream analysis if it is represented by at least 3 reads or UMI counts in at least 2 samples in the time series, to remove TCRs remaining around the detection limit.

### C. Frequency transform

For each timepoint, we derive the counts (*n*_*s,t*_) of each observed sequence *s* at each timepoint *t* = 1, …, *M* . When calculating the log-transformation, we replace *n*_*s,t*_ = 0 by a pseudo-count 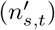 of 1/3 and calculate the frequency 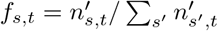 by dividing the counts by the total UMI count from the sample, including the pseudocounts. When calculating the *g*(*f* ) transformation, *f*_*s,t*_ is simply defined as *n*_*s,t*_*/N*_*r,t*_ where *N*_*r,t*_ is the sequencing depth for sample *t*.

For each individual in each dataset, we estimate the *β* and *σ* parameters of Eq. 2 by looking at all pairs of available timepoints and calculating the fluctuations (*δf* ^2^) of each TCR clone. We then bin the TCR clones based on the average frequency of each timepoint pair using the deciles of the average frequency distribution and fit the *b ≡β/N*_*r*_ and *σ* parameters from the resulting distribution across all timepoint pairs (Fig 1). The *σ* parameter estimation is bounded [0, 1] and the *b* = *β/N*_*r*_ parameter estimation is bounded [0, *∞*]. After fitting, we imposed *b ≥*1*/* max_*t*_ *N*_*r,t*_. For further analysis we calculate an average *β* = ⟨*b ×N*_*r*_⟩ for each patient to use for all samples. The distributions of *β* and *σ* across all patients in each dataset are shown in Fig S3.

### D. PCA calculation and outlier detection

Before passing trajectory information to the principal component analysis (PCA), the transformed vector of frequencies at all timepoints of each TCR is normalised by its maximum frequency across the time series, so that clones with large and small frequencies can be aligned. For the log-transform, the normalization is as follows:

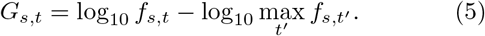

For an equivalent treatment to the log-normalisation, we normalise *g*(*f* ) by subtracting the maximum transformed frequency from each vector:

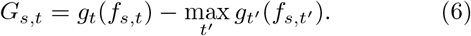

We perform PCA on the centered and normalised *G*,

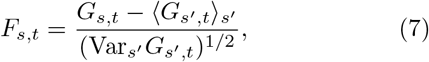

and project the vector **F**_*s*_ = (*F*_*s*,1_, *F*_*s*,2_, …, *F*_*s,M*_ ) onto the principal components:

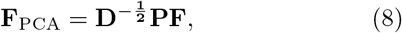

where **D** are the eigenvalues and **P** the eigenvectors of the covariance matrix of **F**_*s*_, 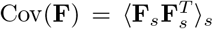. The whitening ensures that the variance is 1 in all principal components (PCs). Practically, PCA is calculated by running `sklearn.decomposition.PCA` with `whiten = True` on the matrix *F*_*s,t*_.

We then use the projection of the frequency vector onto the principal components **F**_**PCA**_ to identify outliers. We assume that the distribution of TCRs in PCA space is dominated by the background sequences that do not expand or shrink, and can be modelled by a normal distribution. By construction of the PCA, its mean is 0 and its covariance matrix is identity, so that:

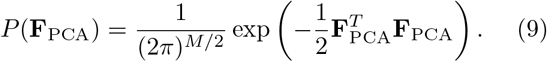

The distribution of distances from the origin, 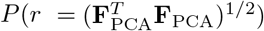 is then given by Eq. 4.

We use the distribution *P* (*r*) to define a threshold *R* to select points that are further from the origin than expected. We allow the user to use a p-value or false-discovery rate (FDR) to select outliers. When using a p-value, we calculate the threshold *R* for which the probability that *r > R* is equal to *p*:

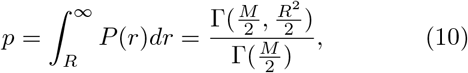

where Γ(*x*) is the Gamma function and Γ(*x, y*) is the upper incomplete gamma function. We use the function `scipy.special.gammainccinv` to solve for *R*.

In the simplest multiple test correction, the p-value is corrected by multiplying it by *S*, where *S* is the total number of unique sequences analysed. Alternatively, we control the false discovery rate (FDR) by computing the number of outliers as a function of *R* in the data, i.e. the number of sequences with *r ≥R, N*_P_(*R*), and compare it to the expected number of outliers we would expect under the null Gaussian hypothesis, which serves as an estimate of the number of false positives 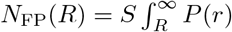 *dr*. The FDR is estimated to be:

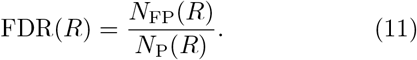

The critical radius *R* is chosen to obtain a desired FDR of 0.05 by inverting that equation.

### E. Clustering of identified outliers

The identified outliers are clustered by calculating the correlation distance between the PCA projections of their transformed frequency trajectories (`scipy.spatial.distance.pdist`), defined as:

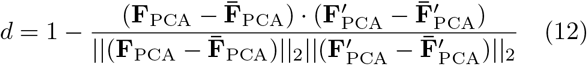

where 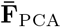 is the mean of the elements of vector **F**_PCA_. Average linkage (`scipy.cluster.hierarchy.linkage`) is then used to create a dendrogram. Clusters are defined by manually selecting a threshold with `scipy.cluster.hierarchy.fcluster`. The sequences from each identified cluster are then plotted on the timeline to identify clusters of interest.

### F. CloneSearch with a subset of timepoints or a lower sequencing depth

To analyse the impact of sample selection on Clone-Search, we took all combinations of 2 to 10 samples. To analyse the impact of average sequencing depth (*N*_*r*_) we down-sampled each sample in the full time series to fractions ranging from 10% to 95% of its sequencing depth in increases of 5%. We repeated the down-sampling 10 times for each fraction.

For parameter estimation, we performed clone filtering on each subsample separately. For combinations of 2 or 3 samples, we included clones that had counts *n*_*i*_ *>* 3 in at least one sample. For combinations of *>* 3 samples, we included clones that had count *n*_*i*_ *>* 2 in at least 2 timepoints. We then calculated the *δf* ^2^ vs *f* profile and fit a *β* and a *σ* for each subsample.

For identification of outliers, we filtered clones and estimated noise parameters in two ways. First, we filtered clones and estimated parameters on each subsample separately, to mimic a real-life situation where a user is handling new samples. Second, we estimated the noise parameters on the complete time series. This mimics a situation where the user inputs parameters from a previously-estimated noise model. For sample subsets, clone filtering was performed on the full time series, retaining clones with count *n*_*i*_ *>* 2 in at least 2 timepoints across the full time series. For *N*_*r*_ down-sampling, clone filtering was performed on each subsample separately. We then calculated outliers by using the *g*(*f* ) transformed frequencies for all TCR clones that passed filtering, using the *β* and *σ* parameters estimated for the whole time series.

### G. Characterisation of outlier properties

For each individual in each cohort, a background set was generated by taking *B* sequences from the nucleotide TCR sequences that passed quality control for that sample, excluding the outliers identified by CloneSearch, where *B* is the number of outliers for that individual.

The probability of a TCR to be observed in a functional repertoire (*P*_post_) [86] was calculated using soN-Nia [87]. To account for differences in sequencing platforms and donor HLA, which we expect to influence selection patterns, we re-fit the selection model to each individual’s full TCR repertoire using the complete list of translated amino acid sequences from all timepoints, including V and J genes, using the training function of soN-Nia. The fitted model for each individual was then used to predict *P*_post_ for the outlier and background TCRs.

## H. Code availability

CloneSearch is available as a Python package at https://github.com/mm523/CloneSearch.

## Acknowledgements

This work utilized resources from the High Performance Computing Group at Memorial Sloan Kettering Cancer Center. M.M. was partially supported by an American-Italian Cancer Foundation Post-Doctoral Research Fellowship. A.M.W. and T.M. were supported by FRM grant EQU202503019997 and ERC PoC grant no. 101185627. This work was also supported by The Olayan Charitable Foundation, The Tow Foundation, The Marcus Foundation, the Ben and Rose Cole Charitable PRIA Foundation (to V.P.B.), the Center for Experimental Therapeutics at MSK (to V.P.B. and B.D.G.) and the Mark Foundation ASPIRE Award (to B.D.G.).

## Author contributions

M.M., Z.S., Y.E., V.P.B., B.D.G., A.M.W. and T.M. conceived the study and developed the methodology, with contributions from all authors. M.M., Z.S., C.R., V.P.B., A.M.W. and T.M. curated the data. M.M. carried out the analyses with contributions from Z.S., S.M., Y.E., V.P.B., B.D.G., A.M.W. and T.M.. All authors interpreted the results. M.M., A.M.W. and T.M. wrote the first version of the manuscript, and all authors reviewed and edited it.

---

**FIG. S1.**
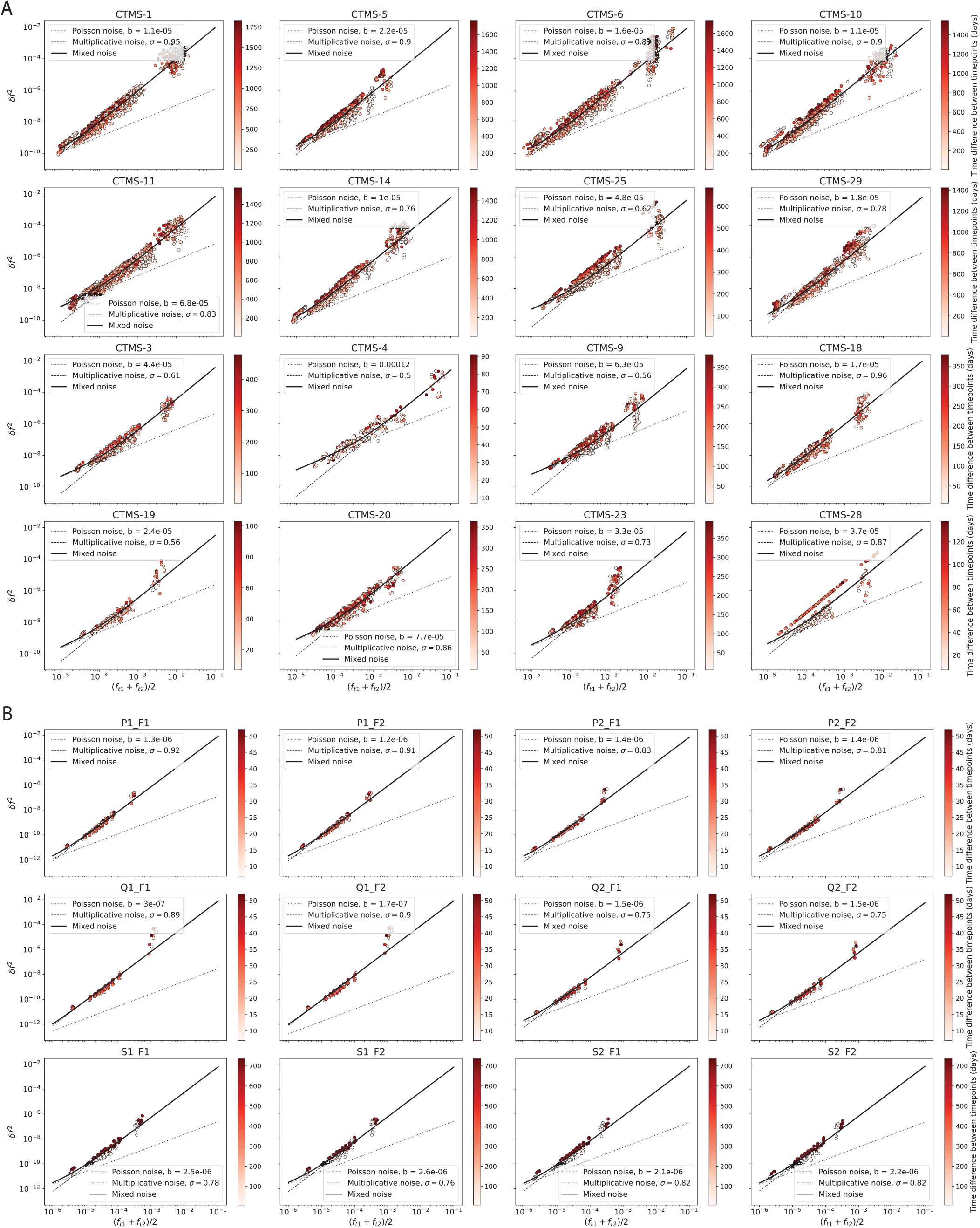
Noise profiles in vaccination datasets. Mean squared difference between timepoints (*δf* ^2^) of TCRs in each individual in the (**A**) PDAC-Vax and (**B**) YFV-Vax cohorts, grouped by frequency as in Fig. 1c.

**FIG. S2.**
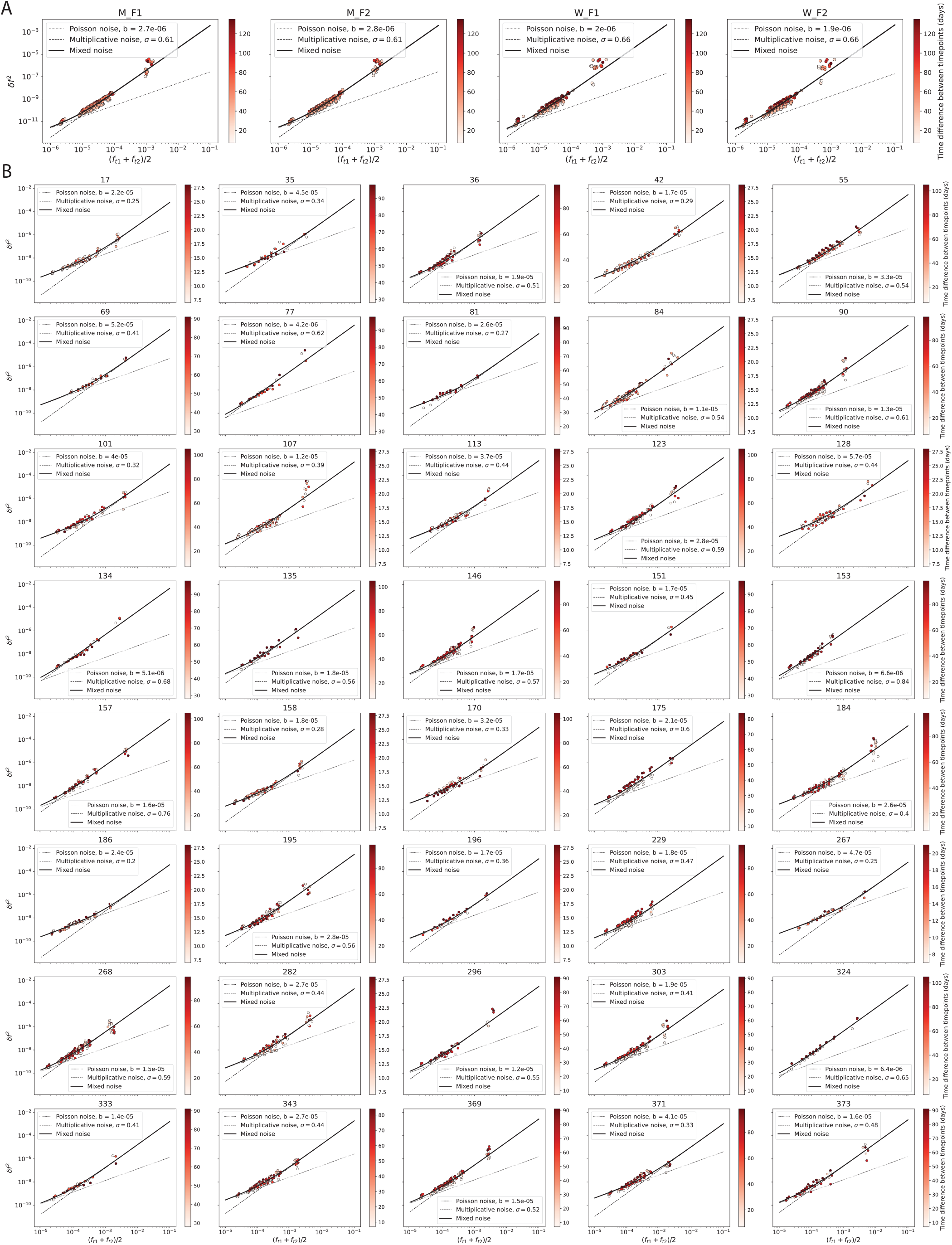
Noise profiles in COVID datasets. Mean squared difference between timepoints (*δf* ^2^) of TCRs in each individual in the (**a**) Minervina-COVID and (**b**) COVIDsortium cohorts, grouped by frequency as in Fig. 1c.

**FIG. S3.**
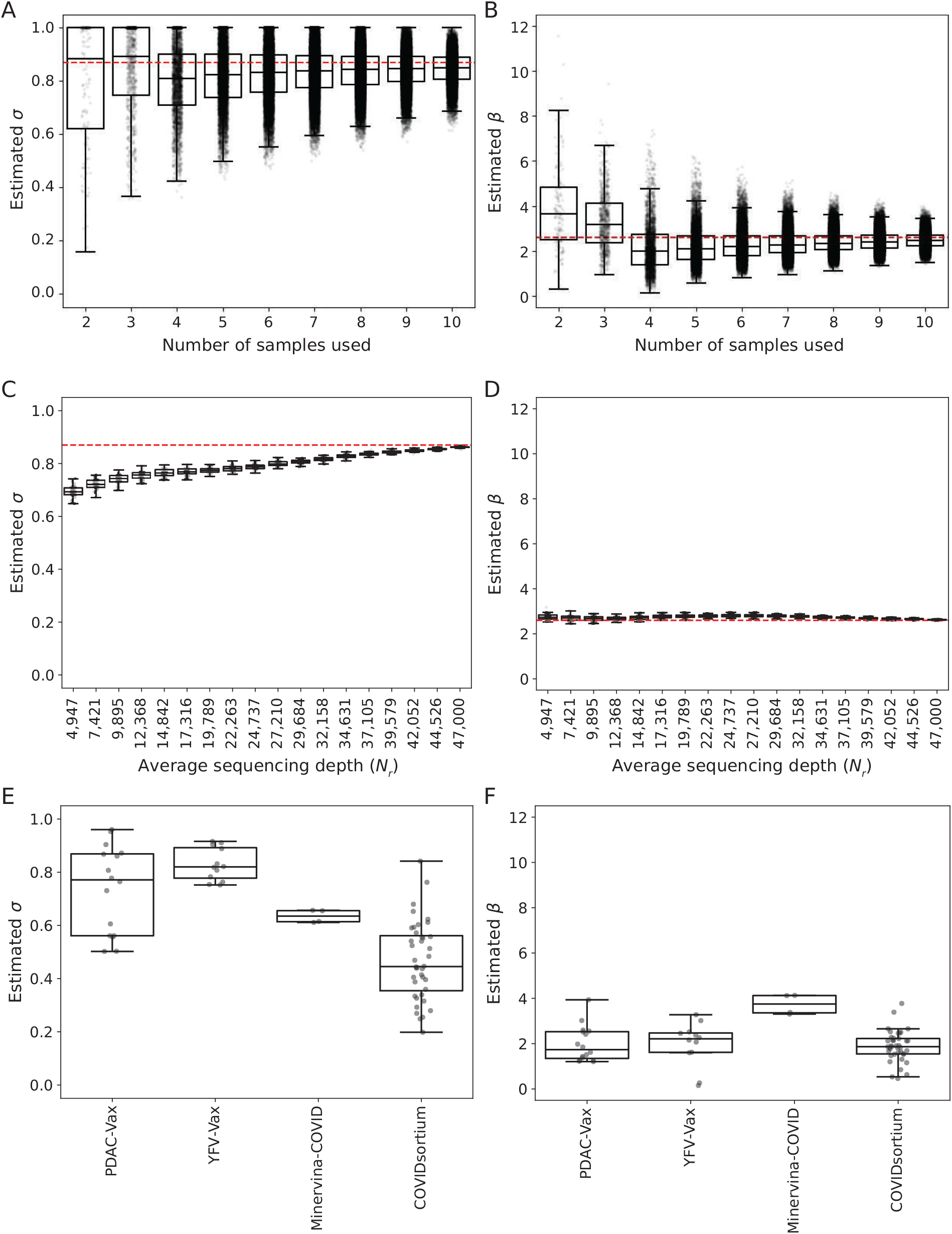
Estimation of *σ* and *β* parameters for varying sample numbers, sequencing depth and different cohorts. **A-B)** Estimated *σ* (**A**) and *β* (**B**) parameters for different subsets of samples from the PDAC-Vax patient 11 time series. All subsets for each size are used. The dashed line indicates the value from the full time series (*σ* = 0.87, *β* = 2.6). **C-D)** Estimated *σ* (**C**) and *β* (**D**) parameters for different average *N*_*r*_ of samples from the PDAC-Vax patient 11 time series. Each sample in the time series is subsampled 10 times to a fraction of its original *N*_*r*_ . The dashed line indicates the value from the full time series (*σ* = 0.87, *β* = 2.6). **E-F)** *σ* (**E**) and *β* (**F**) parameters estimated from the noise profiles of each individual time series in each cohort.

**FIG. S4.**
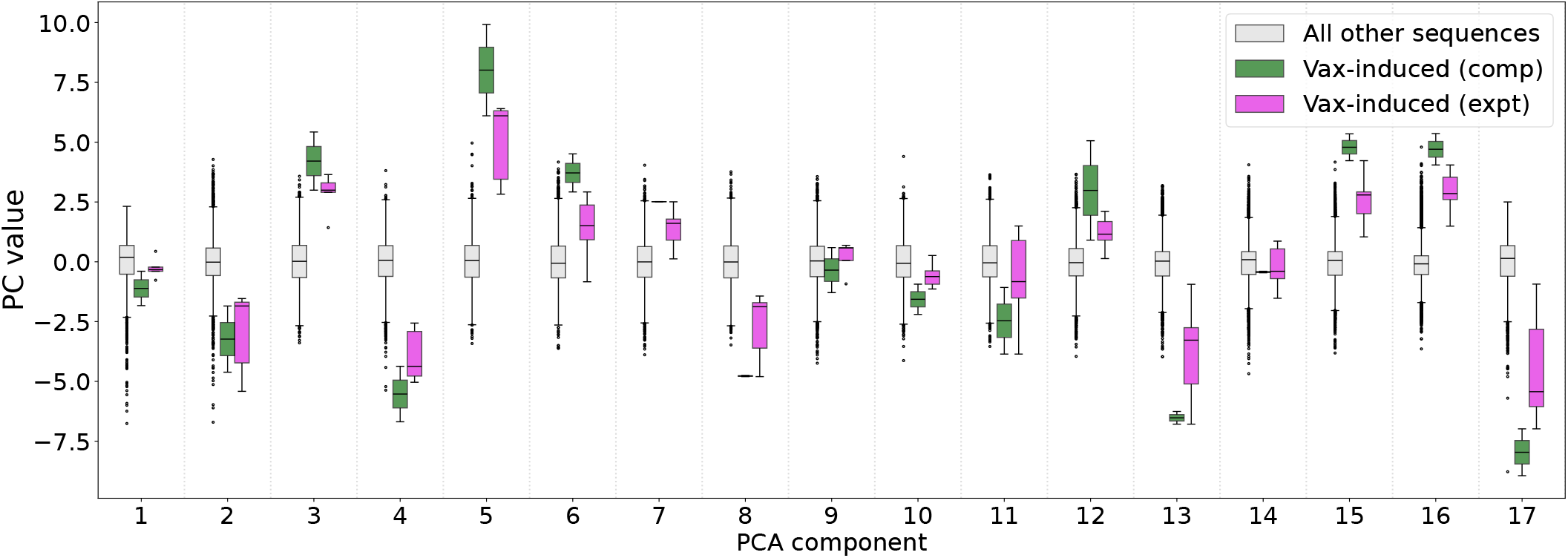
Distribution of annotated and background TCRs on all principal components. Example PCA on PDAC-Vax patient 11. Boxplots show the distribution of all (grey) and previously annotated clones (green and pink).

**FIG. S5.**
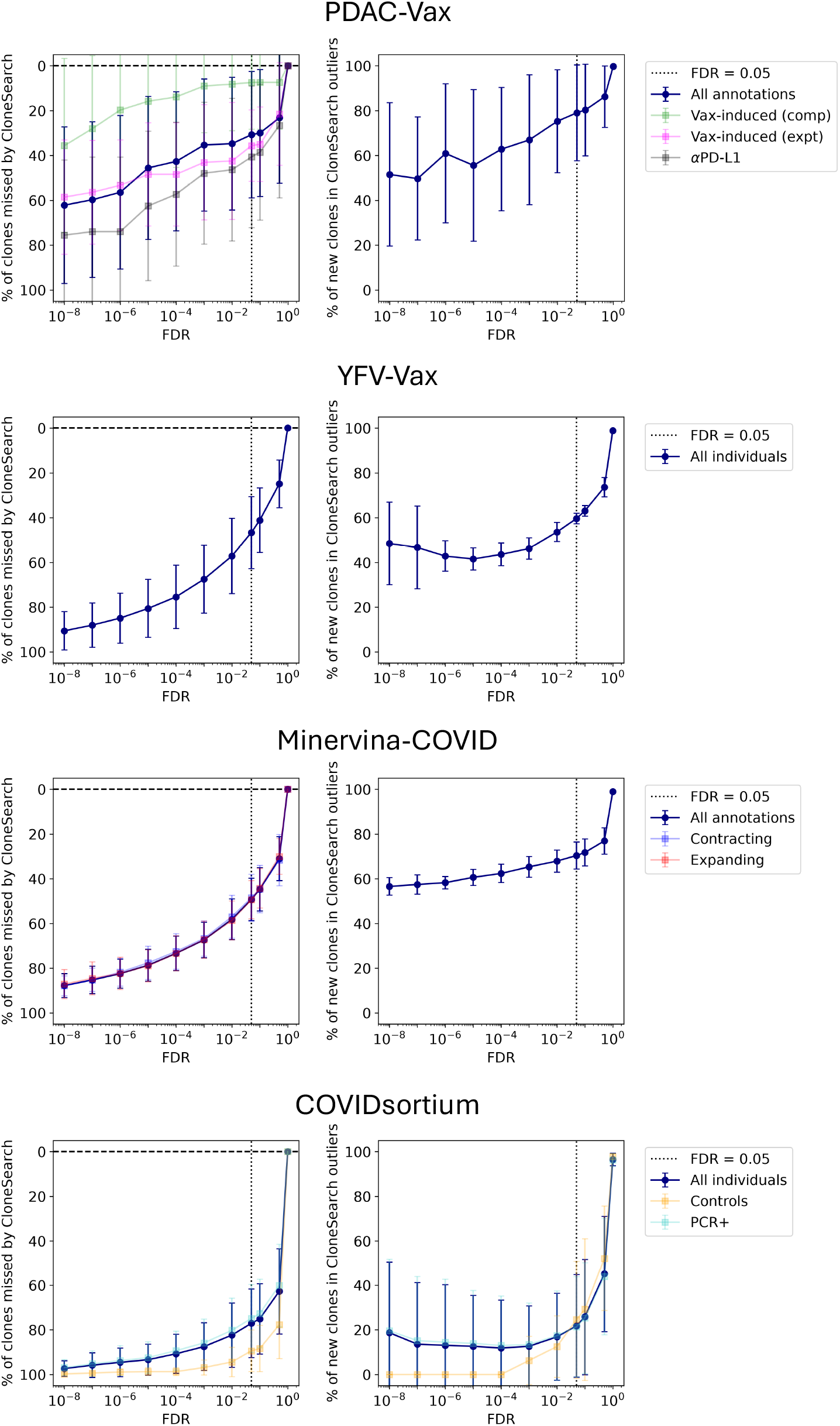
Identification of published sequences and new clones by varying FDR thresholds. For each cohort, the average percentage of previously annotated clones identified by CloneSearch is plotted on the left panel. On the right panel, the percentage of CloneSearch clones that does not overlap with known annotated sequences is shown. The calculations are performed per-patient and the mean calculated across all patients is shown. The error bars show the standard deviation (SD) across all patients. Patients with no outliers for a specific FDR or with no annotated clones of a specific category are removed from the mean and SD calculation.

**FIG. S6.**
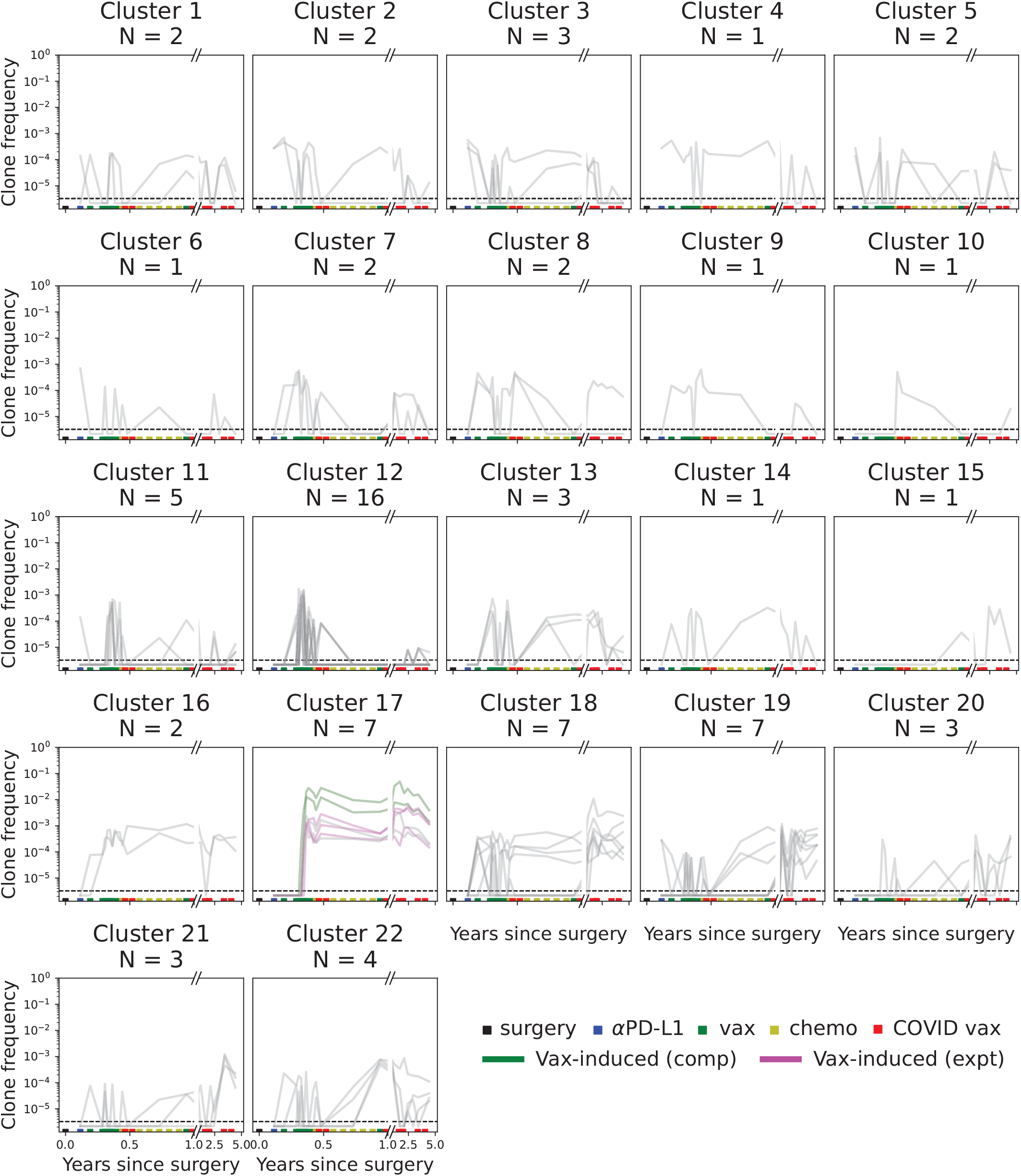
Dynamics of outliers in PDAC-Vax patient 11. Trajectories of all 76 clones identified for PDAC-Vax patient 11 by PCA, organised in clusters as in Fig 2e.

**FIG. S7.**
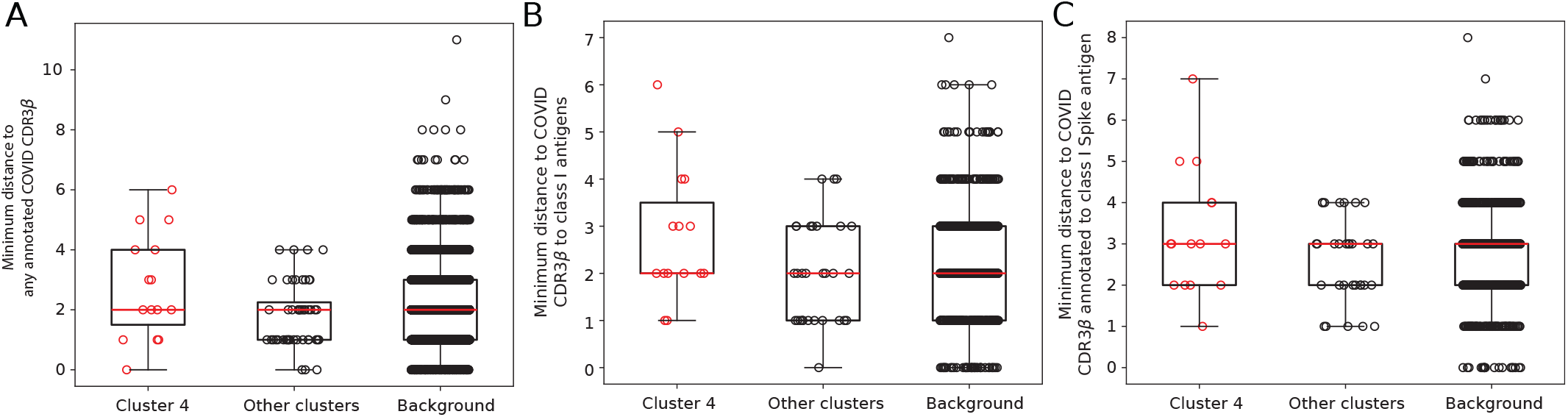
Similarity of putative COVID TCRs to previously annotated COVID-specific TCRs. To validate the putative COVID response identified, we downloaded previously annotated COVID reactive CDR3*β* sequences from 3 publicly available datasets [94–96]. We then calculated the minimum edit distance of each CDR3*β* amino acid sequence in either the putative COVID cluster (cluster 4), the other clusters of outlier TCR clones, or all clones that pass the detection threshold to any CDR3*β* in these sets of known COVID-specific CDR3*β* sequences (**A**). We did not find an enrichment for CDR3s with higher similarity to previously annotated COVID sequences in this cluster, compared to outliers in other clusters or compared to the other TCR clones that pass the detection threshold in this patient. We then refined the search to specifically CDR3*β* sequences annotated to class I epitopes from [95, 96] (**B**), since the putative response identified comprises only CD8 T cells (Fig 4c). Finally, since we hypothesise that these TCRs are a result of vaccination, we further restricted the similarity search to sequences annotated as specific to the Spike protein (**C**). None of the comparisons showed an enrichment of previously annotated COVID sequences in identified outliers.

**FIG. S8.**
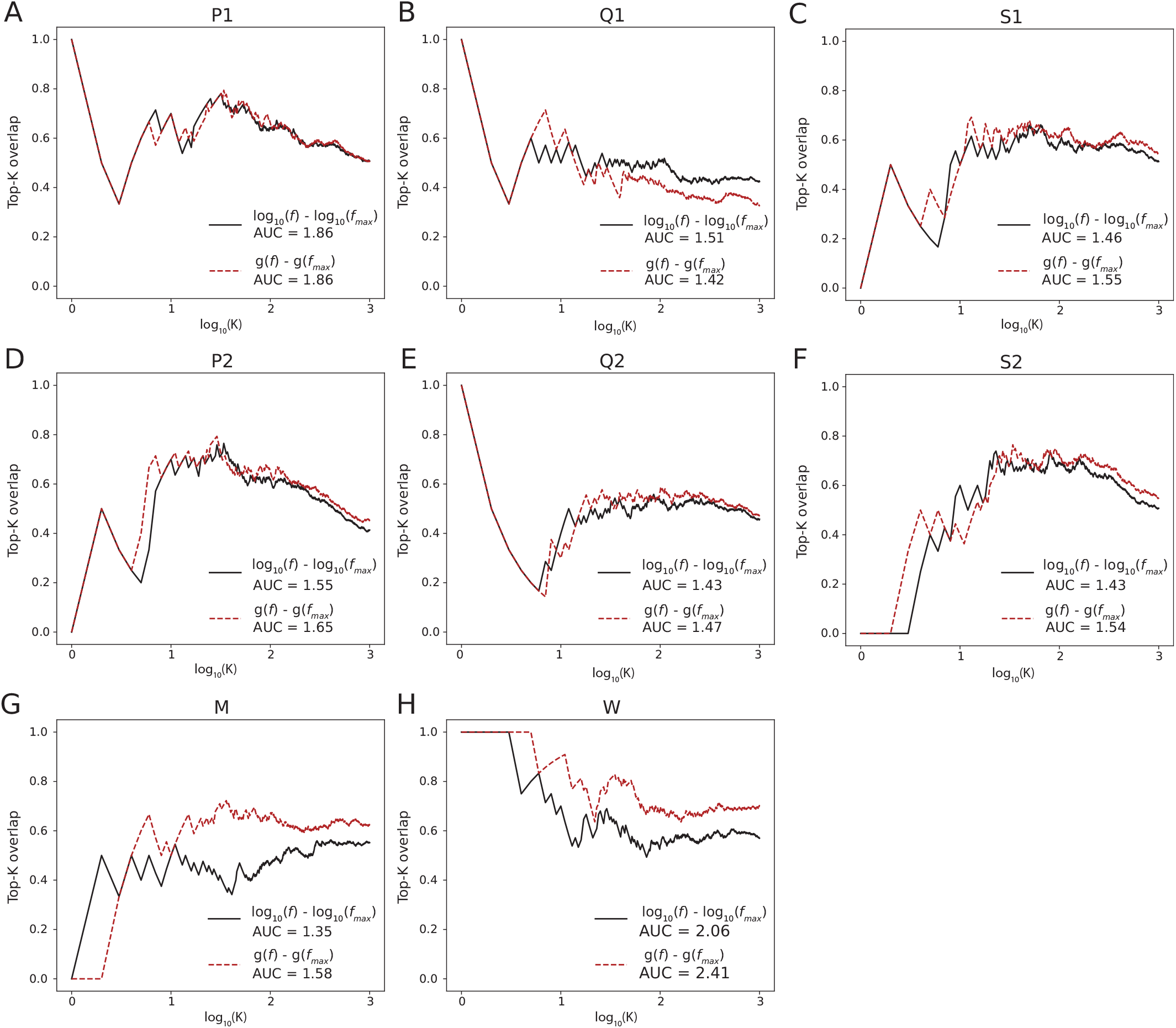
Top-K overlap curves between sequencing replicates. Proportion of shared outliers between two replicate time series for the same patient in the (**A-F**) YFV-Vax and (**G-H**) Minervina-COVID cohorts. The overlap is calculated as the intersection of the top-*K* clones for replicate F1 and F2, divided by *K*. To compare across individuals, we measure the area under these curves.

## A Appendix

### 1. Importance of sample selection in longitudinal analyses

Leveraging the dense sampling from PDAC-Vax patient 11, we created time series with all possible combinations of 2 to 10 samples from the original dataset as in Fig S3a, b. For each combination of samples, we performed outlier selection with CloneSearch (see details in Methods), estimating *β* and *σ* from each smaller time series and compared the results to outliers found using all samples (“full-series outliers”, Fig A1a, b; FDR = 0.05 in all cases). As expected, more of the full-series outliers are identified when more samples are included (Fig A1b). Since parameter estimation in these reduced time series is variable (Fig S3a, b), to mitigate the effects of the difference in parameter estimation and evaluate more cleanly the impact of sample selection, we also ran Clone-Search on each sample subset using *β* = 2.6 and *σ* = 0.87, as calculated from the full time series (Fig 1c). Overall, we find that the number of identified outliers is stable regardless of the number of samples included, but with large variability depending on the sample selection (Fig A1c). In this setting too, we find that including more samples increases the proportion of full-series outliers identified, although with a large spread (Fig A1d). The observed variability in the number of identified outliers and percentage of full-series outliers identified by different subsets of samples is consistent with our hypothesis that outlier identification depends on sample selection: certain timepoint combinations may not capture the relevant trajectory information that would distinguish full-series outliers from the background. To evaluate this, we looked at which combinations of 3 samples best recapitulate previously identified vaccine-induced clones for this patient (“Vax-induced (comp)” and “Vax-induced (expt)”, as defined in Fig 2). We find that the most successful subsamples include: (1) a sample from pre-treatment (leftmost 2 samples in the time series) or early timepoint in the vaccination, when vaccine-induced clones are at low frequency or undetectable (Fig 2f); (2) a sample from either the vaccination timepoints or the chemotherapy timepoints, when frequency of vaccine-induced T cell clones is maximal; and (3) a sample from the follow-up timepoints (Fig A1e). In summary, the outlier detection approach is not very sensitive to the exact choice or number of timepoints used in the analysis, as long as they are in sufficient numbers and correctly frame the time of immunological scarring of interest.

We then study how outlier selection performs when sequencing depth (*N*_*r*_) is reduced by systematically subsampling an existing time series, and asking how outlier identification changes. For the PDAC-Vax patient 11 time series, we subsampled each sample in the time series to a fraction (0.1 to 0.95) of its initial *N*_*r*_ 10 times and performed outlier selection. As above, we performed outlier detection both by using subsample-specific *σ* and *β*, and using *β* = 2.6 and *σ* = 0.87 (as estimated from the full time series) for all subsamples, to limit the impact of noise estimation on the selection process (Fig A2). The number of identified outliers in each subsample depends heavily on sequencing depth: the smaller the sample size, the fewer outliers are detected (Fig A2a). This is the expected behaviour, as lower sequencing depth will result in fewer clones, and therefore a smaller proportion of clones that can be detected as outliers. In line with the fewer overall outliers, the percentage of full-depth outliers (defined as responding TCRs identified by CloneSearch with no subsampling) identified in each subsample also depends strongly on the average sequencing depth (Fig A2d, black dots and boxes). However, our use of a minimum detection threshold for inclusion of clones in the analysis (see Section A 2) introduces a confounding factor: with lower sequencing depth, fewer TCRs will pass the threshold, and so fewer of the full-depth outliers will be available for the method to detect (Fig A2c). When correcting for the number of full-depth outliers that pass the clone detection threshold, the percentage of recovered full-depth outliers increases (Fig A2d, red dots and boxes; Fig A2e). The results from using subsample-specific noise parameters or shared noise parameters across all subsamples are consistent, although more outliers are identified at smaller average sequencing depths when using subsample-specific parameters (Fig A2a, b). However, the larger number of outliers does not result in better capture of full-depth outliers (Fig A2e, f). This observation is likely a consequence of the lower *σ* values estimated for subsamples with smaller average sequencing depths (Fig S3c), which underestimates the noise, and overestimates the fluctuations in background clones.

### 2. Selection of clones for inclusion in longitudinal analysis

We wondered whether CloneSearch could provide a more principled way of performing the clone filtering, in a way that can be tuned for any time series. To this end, for each clone we calculate its total count across all timepoints. We then filter clones using a threshold *C* on the summed counts. At *C* = 0, all clones that are seen in at least one sample are included. At *C* = 2, only clones with a count of 2 in at least one sample, or clones which are observed in at least 2 samples are included, and similarly for larger *C*. We calculate the PCA projection of clones that pass filtering at each *C*, for *C* that varies from 0 to 15 and *C* = 20, to observe how the distribution of clones in PCA space changes. We find that the *P* (*r*) distribution from the PCA projection is jagged at *C* = 0 due to the number of small clones that are detected at only one timepoint, but becomes smoother as *C* increases (Fig A3A, example shown for PDAC-Vax patient 11). At higher *C*, the observed distribution of radii in PCA space becomes more similar to the expected radial distribution under the Gaussian null model (Fig A3A). To find the optimal *C* for a time series, we calculate the Kolmogorov-Smirnov D-statistic, which calculates the difference between the observed and assumed cumulative distribution functions of *P* (*r*) (Fig A3B). For PDAC-Vax patient 11, the D-statistic is minimal at *C* = 11. We then looked at how the set of identified outliers changes as *C* increases, to evaluate whether stricter filtering changes the set of called outlier clones, or simply reduces the set to a cleaner subset. Performing outlier identification with CloneSearch using the adaptively-tuned *C* = 11 reduces the number of observed outliers compared to lower *C* thresholds, most of which are removed because of their *C* (Fig A3C), but does not change the relative radii of the maintained set (Fig A3D). With this method, 64 of 76 outliers found in Fig 2 are still identified, and an extra 31 new outliers are added to the set. Interestingly, these newly-identified outliers are concentrated within two specific clusters: the previously-identified CD4 cluster (cluster 9 in Fig A3E), and a newly observed cluster with clones that are solely detected at timepoint chemo-1. Thus, projection of TCR clones in PCA space may provide a principled way to filter clones for downstream analysis. We provide this tunable clone filtering strategy as an option within CloneSearch.

**FIG. A1.**
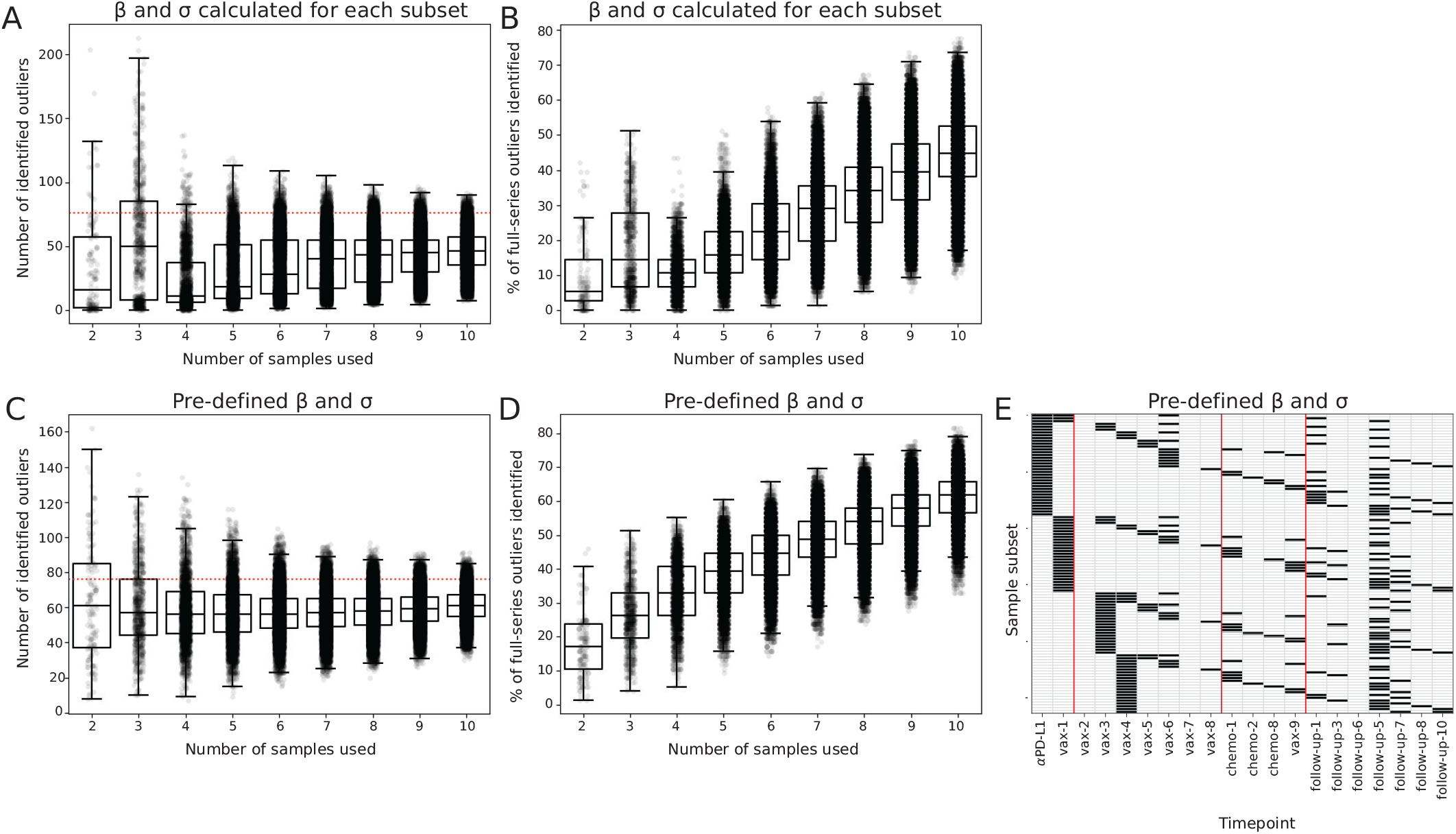
Impact of number of sequenced samples and sample selection on outlier selection, for PDAC-Vax patient 11. **A-B)** Impact of including fewer samples on outlier selection, quantified as the total number of identified outliers (**A**) and the percentage of full-series outliers identified when fewer samples are used (**B**). We calculated the percentage of outliers identified by using the full time series for PDAC-Vax patient 11 that are still called as outliers by taking fewer samples. *β* and *σ* parameters are estimated for each subset of samples separately. **C-D)** Impact of including fewer samples on outlier selection, quantified as the total number of identified outliers (**C**) and the percentage of full-series outliers identified when fewer samples are used (**D**), when the parameter estimation and the clone filtering are performed on the full time series (*β* = 2.6 and *σ* = 0.87). **E)** Sample selection remains important to identify certain categories of outliers. We looked at the subsets of 3 samples that allowed us to identify *>* 70% of previously identified vaccine-induced clones. Each row is a subsample and black boxes indicate which timepoints are included.

**FIG. A2.**
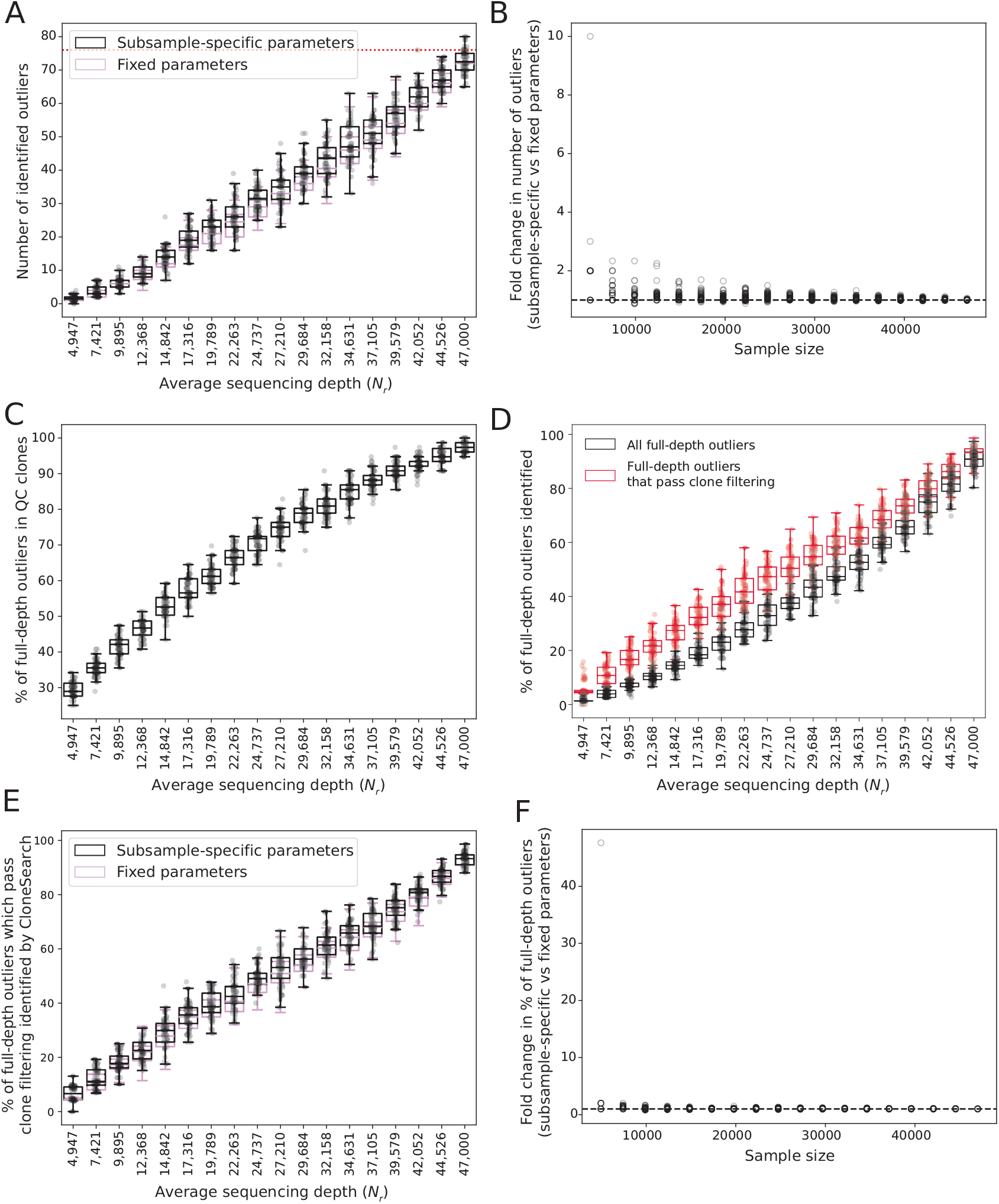
Impact of sequencing depth on outlier selection in PDAC-Vax patient 11. **A)** Number of outliers detected when each sample in the time series is subsampled to a fraction of the starting *N*_*r*_ (10%-95%) to simulate lower sequencing depth, plotted as a function of the average read count *N*_*r*_ across all samples. The outlier identification is performed by using noise parameters estimated for each subsample (black), or fixed parameters estimated from the complete time series (purple, *β* = 2.6 and *σ* = 0.87) when subsampling *N*_*r*_ . **B)** Ratio of the number of outliers detected by using subsample-specific noise parameters and fixed parameters when subsampling *N*_*r*_, defined as in **A. C)** Relationship between average sequencing depth (*N*_*r*_ ) and the number of outliers obtained from the full time series that pass clone filtering (see Methods Section VII F). Outliers are identified using fixed parameters from the full time series. **D)** Percentage of full-depth outliers, defined as outliers detected when all reads are used, re-captured in each *N*_*r*_ subsample (black). Since many full-depth outliers do not pass the clone filtering threshold following subsampling (**C**), we also report the proportion among those that did pass the detection threshold (in red). Both are run using fixed parameters from the full time series. **E)** Percentage of the full-depth outliers that pass clone quality control identified by CloneSearch when using subsample-specific noise parameters (black) or fixed parameters (purple) in *N*_*r*_ downsampling. **F)** Ratio of the percentage of the full-depth outliers that pass quality control identified by CloneSearch when using subsample-specific noise parameters and fixed parameters in *N*_*r*_ downsampling, defined as in **E**.

**FIG. A3.**
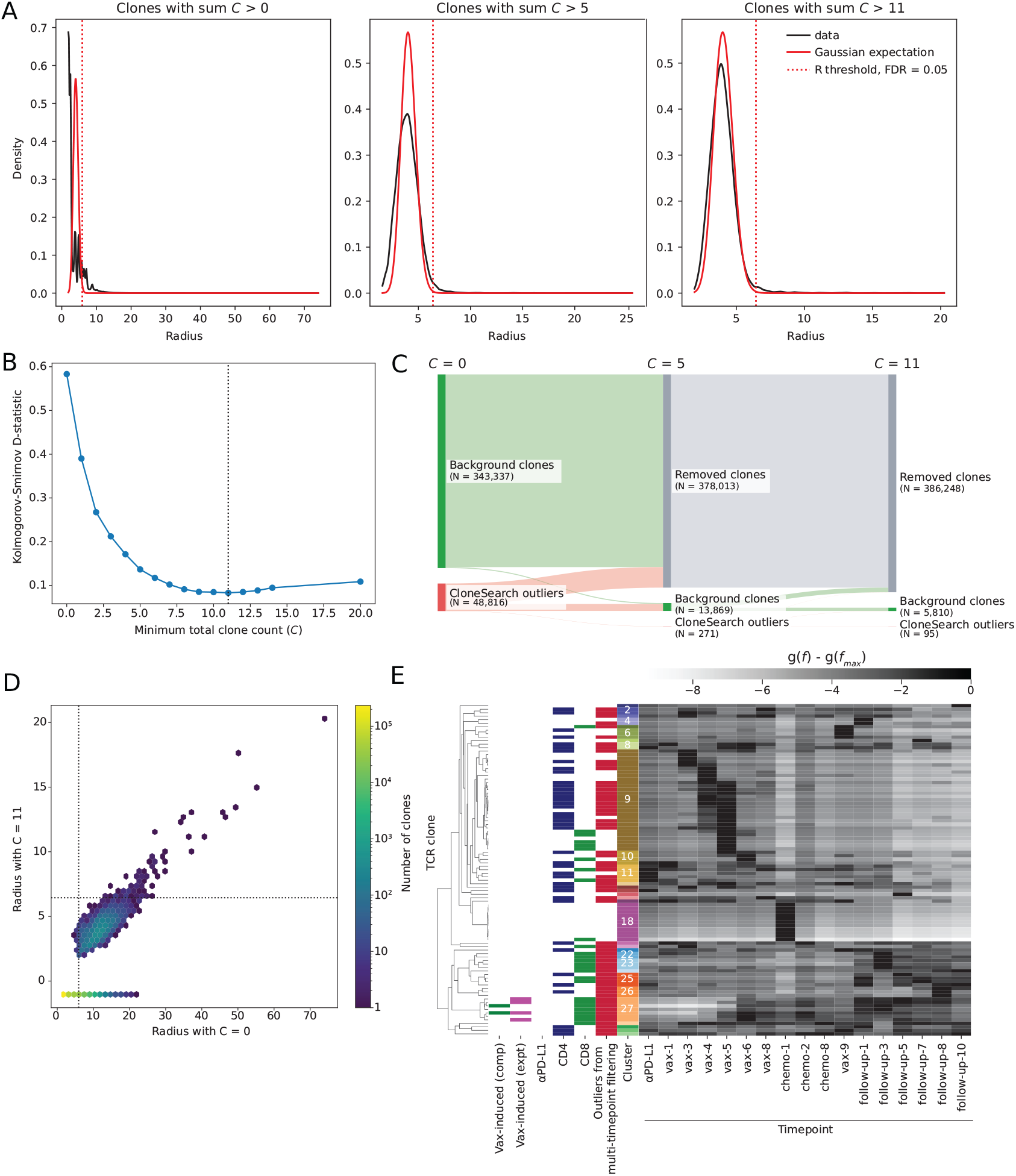
Tunable selection of clones to include in the CloneSearch analysis. **A)** Distribution of observed (black) and expected radii (*P* (*r*)) in PCA space when different thresholds *C* are used to select clones from the time series. *C* represents the total counts for a clone across all samples. **B)** Kolmogorov-Smirnov goodness-of-fit test statistic of observed and expected *P* (*r*) when *C* is increased from 0 to 20. **C)** Sankey plot to represent the fate of clones that are included when *C* = 0 when *C* is increased to 5 (middle column) or 11 (right-most column). In grey, green and red, clones that are excluded, identified as background or identified as outliers for each *C*. **D)** Correlation of the radii for all clones when *C* = 0 and *C* = 11. Clones that are removed at *C* = 11 but present at *C* = 0 are plotted with radius *<* 0 on the y-axis. The colour of each hexagon represents the number of clones in that hexagon. **E)** Heatmap of identified outliers with *C* = 11. The same annotations as in Fig 4A are provided. The red column (“Outliers from multi-timepoint filtering”) represents which of these outliers are also found in Fig 4A. Clustering was performed with average linkage and correlation distance, clusters were defined at a distance threshold of 0.3.

